# A small RNA guides a post-transcriptional regulatory protein to its target

**DOI:** 10.64898/2026.08.12.744399

**Authors:** Raya Faigenbaum-Romm, Lior Aroeti, Netanel Elbaz, Shira Fisher, Yael Altuvia, Liron Argaman, Meshi Barsheshet, Miriam Ravins, Naama Katsowich, Yifei Xiong, Sigal Ben-Yehuda, Hanah Margalit, Ilan Rosenshine

**Affiliations:** Department of Microbiology and Molecular Genetics, Institute for Medical Research Israel-Canada, Faculty of Medicine, The Hebrew University of Jerusalem, Jerusalem 9112102, Israel; Racah Institute of Physics, The Hebrew University of Jerusalem, Jerusalem, Israel

## Abstract

RNA-binding proteins are central to post-transcriptional regulation and are generally thought to recognize their target RNAs through direct binding. Here, we uncover a previously unrecognized mechanism in which a small RNA (sRNA) guides a protein regulator to specific target mRNAs, enabling programmable post-transcriptional control. By mapping in pathogenic *Escherichia coli* the RNA interactome of CsrA, a global post-transcriptional regulator, we identify ∼800 ternary complexes in which CsrA simultaneously binds an sRNA and an mRNA. Focusing on one class of these complexes, we show that PasE, a newly discovered sRNA, directs CsrA to virulence-associated mRNAs that are otherwise not targeted by CsrA, resulting in their repression. Altering the PasE seed sequence redirects CsrA to selected mRNAs, establishing a modular RNA-guided platform for gene regulation. Together, our findings reveal a new and likely widespread mechanism of bacterial post-transcriptional regulation and provide a framework for a new type of RNA-based synthetic regulation.

**Highlights:**

- The PasE sRNA can direct the CsrA post-transcriptional regulator to target mRNAs that are otherwise not targeted by CsrA.
- The CsrA-PasE complex controls the virulence of enteropathogenic *E. coli*.
- The PasE seed can be synthetically reprogrammed to direct CsrA to new target mRNAs.
- Hundreds of additional sRNA-CsrA-mRNA regulatory complexes are formed in pathogenic *E. coli*.

## Introduction

The Carbon Storage Regulator A (CsrA) is a widely conserved bacterial RNA-binding protein that controls key processes, including biofilm formation, motility, carbon metabolism, and virulence^1^. In *E. coli*, CsrA binds to hundreds of mRNAs, influencing their stability or translation^2^. It functions as a homodimer, with each subunit binding to a GGA motif within RNA hairpin loops, altering RNA topology to expose or block functional regions, such as ribosome or Rho binding sites^1, 3^. CsrA activity is sequestered in *Escherichia coli* by the small RNAs (sRNAs) CsrB and CsrC, which contain multiple CsrA-binding sites^4, 5^. Additional sRNAs, including Spf, McaS, and GadY, harbor one or two CsrA-binding sites and may similarly modulate its availability^2, 6–9^. Several of these sRNAs are incorporated into CsrA condensates, suggesting an additional layer of spatial regulation^10^. In pathogenic *E. coli* strains, such as enteropathogenic *E. coli* (EPEC), CsrA serves as a central post-transcriptional regulator of virulence, linking metabolic state to infection capacity^10–13^.

EPEC causes acute intestinal lesions and chronic diarrhea, posing a major health risk to young children^14, 15^. Its virulence relies primarily on two surface machineries: The Type III Secretion System (T3SS), a syringe-like apparatus that injects effectors into host cells, and the type IV Bundle-Forming Pili (BFP), which promote host attachment. The T3SS genes are clustered within the chromosomal Locus of Enterocyte Effacement (LEE)^16, 17^, whereas the BFP genes are encoded on the pMAR2 plasmid, including the recently discovered *pilW* gene^15, 18–20^. To ensure successful host colonization, the expression of T3SS and BFP is co-regulated via multiple transcriptional and post-transcriptional factors^21–23^. The primary post-transcriptional regulator of EPEC virulence is CsrA, critical for T3SS and BFP expression^10–13^. Notably, CsrA also coordinates the transition of EPEC from planktonic to host-attached lifestyle^11–13, 24^. However, the underlying mechanisms and the global CsrA regulatory network remain poorly understood. To systematically define CsrA-mediated regulation at the RNA level, we used RIL-seq (RNA Interaction by Ligation and Sequencing), a powerful approach for mapping RNA-RNA interactions mediated by an RNA-binding protein, which has been successfully applied to the global RNA-binding regulators Hfq and ProQ^25–27^. Here, we applied RIL-seq to map the CsrA-RNA interactome, uncovering an extensive network of binary (CsrA-RNA) and ternary (RNA-CsrA-RNA) complexes. By investigating the significance of ternary complexes, we discovered that a newly identified sRNA, PasE, modulates EPEC virulence by guiding CsrA to its target mRNA. Given the CsrA conservation across the bacterial domain, we predict that this mechanism is likely widespread.

## Results

### Formation of CsrA-RNA binary and ternary complexes

To elucidate the CsrA regulatory network in EPEC, we applied RIL-seq^25, 26^, a high-throughput method that has enabled us to systematically determine binary and ternary CsrA-RNA complexes (Fig. 1a and Fig. S1). In RIL-seq, RNAs bound to the same protein are coimmunoprecipitated, ligated, and sequenced, to identify chimeric RNAs that map to two distinct genomic locations, representing RNA pairs co-bound to the protein, possibly interacting. RIL-seq libraries were generated from EPEC (strain NN6861) grown under conditions that promote expression of its virulence genes (host-mimicking conditions)^23^, followed by sequencing and analysis via our RIL-seq computational pipeline^25, 26^. The results of six biological replicates are summarized in Supplementary Table 1, 2, and 3, reporting both single and chimeric RNA fragments. RIL-seq identified 1,241 CsrA-bound RNAs, each mapped to a distinct chromosomal location (“singles”; Supplementary Table 3), reflecting CsrA dimers bound to one RNA molecule or two identical ones (Fig. 1a, left panel). Most singles corresponded to the canonical CsrA-sponge sRNAs CsrB (57%) and CsrC (21%) (Fig. 1b)^1, 28^. The remaining singles corresponded mostly to mRNA, including both coding and untranslated regions, and to a lesser extent to tRNAs (Fig. 1b). Importantly, we identified 818 statistically significant chimeric RNA fragments mapped to two different chromosomal locations. These fragments, termed chimeras (aka statistically significant chimeras, or S-chimeras^25, 26^), represent RNA pairs bound to the same CsrA dimer (Supplementary Table 2). These results indicate frequent formation of CsrA-RNA ternary complexes (Fig. 1a right panel). Driven by their potential to reveal new concepts in post-transcriptional regulation and virulence control, we set out to investigate the biological significance of these ternary complexes.

**Figure 1.**
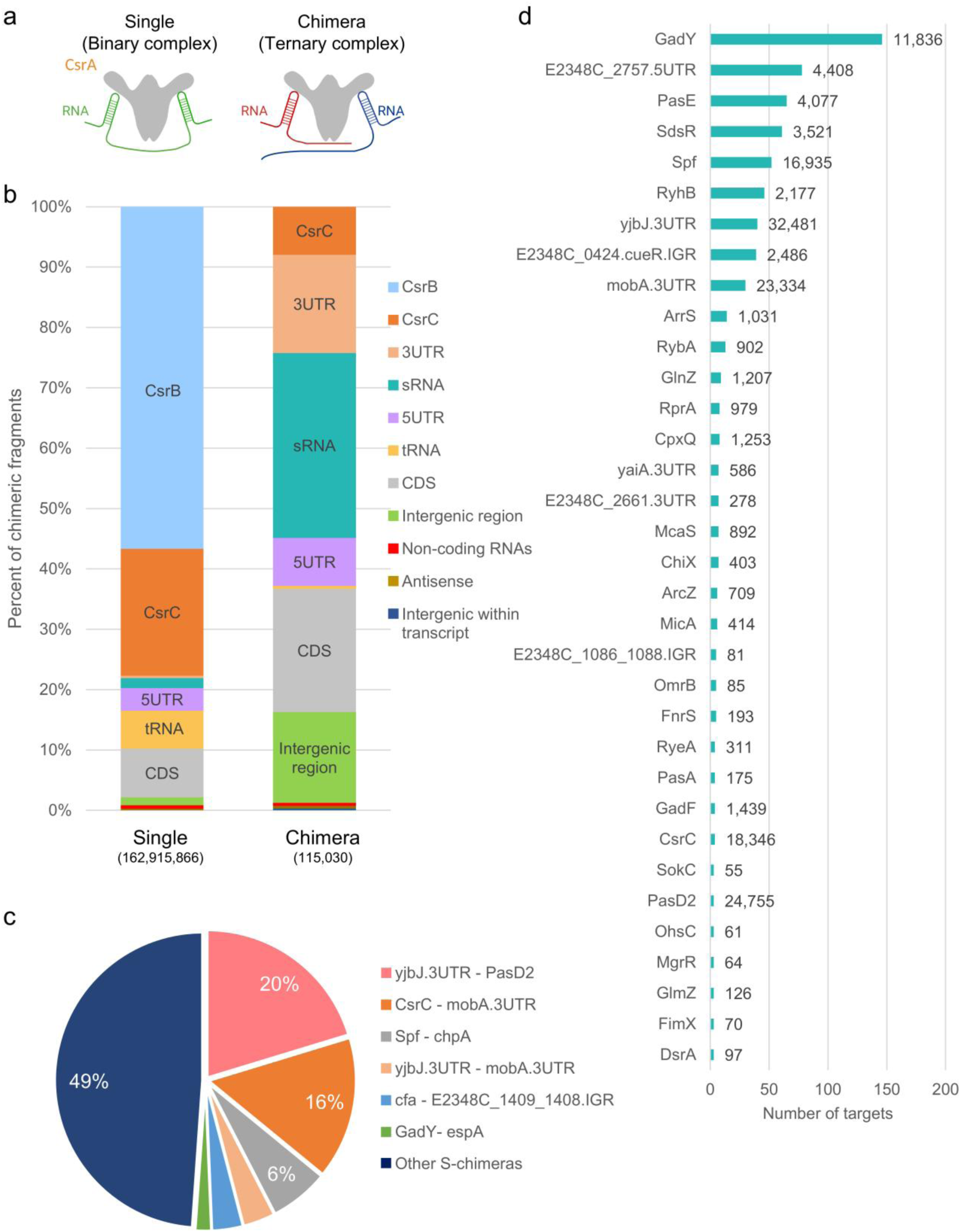
CsrA forms binary and ternary complexes with RNAs. (a) Schematics of CsrA-RNA complexes. Left: A CsrA-RNA binary complex. A CsrA dimer bound to a single mRNA molecule (green) containing two CsrA binding sites, identified as a single RNA in the RIL-seq analysis. Right: A CsrA-RNA ternary complex. CsrA dimer bound to two different RNA molecules (red and blue), identified as a chimera in the RIL-seq analysis. (b) Distribution of CsrA-bound single RNAs (left) and chimeric RNA fragments (right), deduced by RIL-seq analysis (Supplementary Table 2 and 3). The bound RNAs were categorized as 5′ untranslated region (5UTR), coding sequence (CDS), 3′ UTR (3UTR), tRNA, sRNA, antisense (AS), intergenic region (IGR), intergenic within transcript (IGT), CsrB, CsrC, and other non-coding RNA excluding tRNA, rRNA, and sRNA (ncRNA). The color code and fraction (%) of each category are indicated. Total counts for detected single and chimeric RNAs are denoted in parentheses. (c) Dominant RNA pairs in CsrA-associated chimeras. Shown are the most abundant RNA pairs identified in more than 2,000 chimeric fragments, summed over all six RIL-seq libraries (Supplementary Table 2). Pair percentage is calculated as the total number of reads of a given chimeric fragment divided by the total number of reads corresponding to all chimeric fragments. (d) Distribution of sRNAs (known and putative) in chimeric fragments. The number of different interacting RNAs co-bound on CsrA with each sRNA is indicated in the X axis. The total number of reads corresponding to chimeric fragments associated with each sRNA is indicated in black next to the respective bar. Only sRNAs with three or more co-bound RNAs are shown. For clarity, the RNA annotated as E2348C_1086.E2348C_1088.IGR (Supplementary Table 2) is abbreviated as E2348C_1086_1088.IGR.

### High abundance of CsrA-bound ncRNA-ncRNA and sRNA-mRNA pairs

The identified CsrA-RNA ternary complexes predominantly involved sRNAs and mRNAs, including intergenic regions (Fig. 1b). Several RNA pairs were highly enriched in the ternary complexes, with the most abundant being the *yjbJ*.3UTR-PasD2 pair, detected in over 23,000 chimeric fragments, accounting for 20% of all chimera-associated reads (Fig. 1c and 1d). PasD2 is a recently discovered prophage-encoded sRNA^23^, whereas *yjbJ.3UTR* likely represents a chromosomally encoded ncRNA. The second most abundant chimera consists of CsrC and *mobA.3UTR*, the latter encoded on the EU580135 small plasmid. This RNA pair was present in over 18,000 chimeric fragments, comprising 16% of the chimera-associated reads (Fig. 1c). Importantly, we also detected numerous cases of regulatory sRNA-mRNA pairs (Fig. S2c), with interacting partners originating from pathogenicity islands and the core genome (Supplementary Table 2). Collectively, these results reveal that CsrA engages in a rich and diverse network of interactions, extending beyond the classical formation of CsrA-mRNA binary complexes.

To define the architecture of the CsrA-driven sRNA-RNA network, the most repeatable statistically significant sRNA-mRNA chimeras were identified as described^29^. This data was used to generate an interaction network composed of CsrA bound with RNA pairs identified in at least four biological replicates (Fig. 2). In this stringent network, the sRNAs GadY and Spf exhibited the highest number of distinct repeatable chimeras (Supplementary Table 2). A few of these sRNA-mRNA pairs were previously reported to be Hfq-bound, particularly for Spf^23^ (Fig. 2, Supplementary Table 2). Five additional putative sRNAs, which formed repeatable chimeras with multiple mRNAs included *yjbJ*.3UTR, *mobA*.3UTR, E2348C_2757.5UTR, E2348C_0424.cueR.IGR, and E2348C_0271.5UTR (Fig. 2). This analysis highlights a complex sRNA-CsrA-mRNA network, suggesting potential involvement of these interactions in regulatory processes.

**Figure 2.**
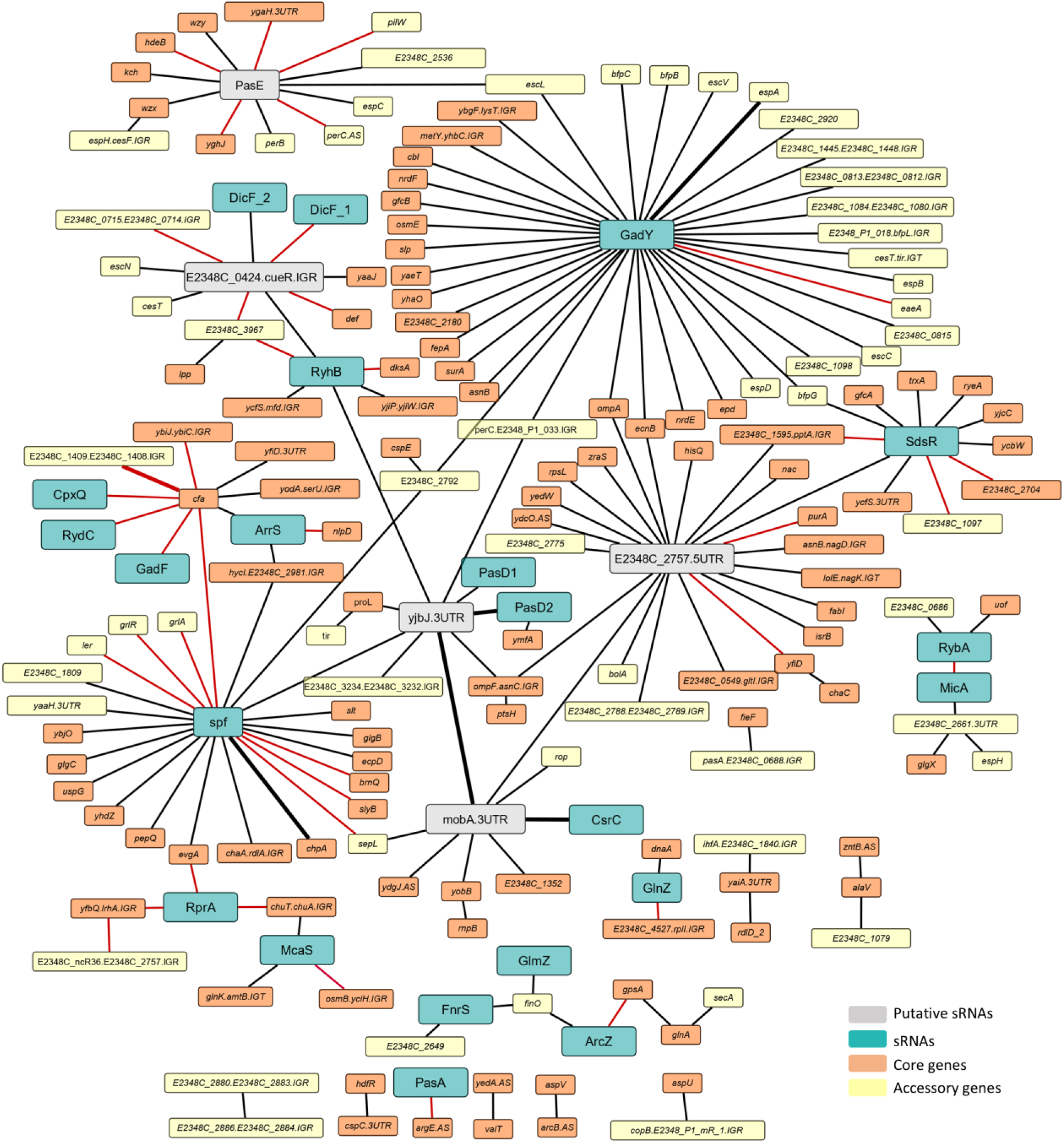
The CsrA-mediated sRNA-RNA interactome. Network representation of the stringent set of chimeras identified in at least four libraries (Supplementary Table 2). Two nodes connected by an edge represent RNAs co-bound to the same CsrA molecule (identified as chimeric fragments). The node colors indicate sRNAs, putative sRNAs, core genes, and accessory genes originating from the LEE, plasmids, prophages, or integrative elements. Black edges represent RNA pairs detected solely on CsrA, and red edges indicate RNA pairs also identified on Hfq ^23^. Thicker edges highlight the most abundant co-bound RNA pairs with at least 2,000 associated chimeric fragments, as shown in Fig. 1c.

### Complementarity between CsrA-bound RNA pairs

Base-pairing potential between the CsrA-bound sRNA and its associated mRNA may further support the involvement of the sRNA in the regulation of the co-bound transcript. To assess this base-pairing potential, we assumed that each set of mRNAs co-bound with a given sRNA might share a common sequence motif that enables base pairing. Using the MEME suite^30^, we identified statistically significant common motifs in mRNAs co-bound with the sRNAs GadY, RyhB, McaS, and the putative sRNAs E2348C_0271.5UTR and E2348C_0424.cueR.IGR (Fig. S3a). Importantly, each of these common motifs was found to be complementary to its corresponding sRNA. Additional groups of potential target mRNAs contained motifs that did not reach statistical significance but nevertheless show complementarity to their respective sRNAs (Fig. S3b). Together, the extensive CsrA-RNA-RNA network shown in Fig. 2 and the sequence complementarity shown in Fig. S3 suggest that the formation of these ternary complexes, many of which include sRNA-mRNA pairs, is biologically meaningful.

### PasE is a CsrA-associated sRNA

To further elucidate the significance of the potential base pairing between the CsrA bound sRNAs and mRNAs, we selected the sRNA E2348C_0271.5UTR for in-depth analysis because it showed high sequence complementarity with all of its co-bound mRNAs (Fig. 2 and Fig. 3a), some of which encode central virulence factors (Fig. 3a). Following previous nomenclature^23^, we renamed E2348C_0271.5UTR PasE (Pathogenicity associated sRNA E) (Fig. S4a). Rapid amplification of cDNA ends (RACE) identified the 5’ and 3’ ends of PasE, defining a 56-nucleotide transcript (Fig. S4b). Northern blot analysis confirmed PasE expression under host-mimicking conditions (exponential growth phase in DMEM at 37 °C) (Fig. S4c-d). Secondary structure prediction revealed that PasE contains two typical CsrA-binding sites and a single-stranded 5’ region that shows sequence complementarity with co-bound mRNAs and thus may function as a seed for target recognition (Fig. 3b-c). PasE, including its putative seed and CsrA-binding sites, is conserved across *E. coli* and *Shigella* species (Fig. S4e, 5 and Supplementary Table S4). Finally, we found that PasE levels were strongly reduced in EPEC deleted of *csrA*, and in EPEC strains containing point mutations in the PasE-CsrA binding sites (Fig. 3d, Fig. S4f, Fig. S6a-b), demonstrating that PasE stability depends on binding to CsrA.

**Figure 3.**
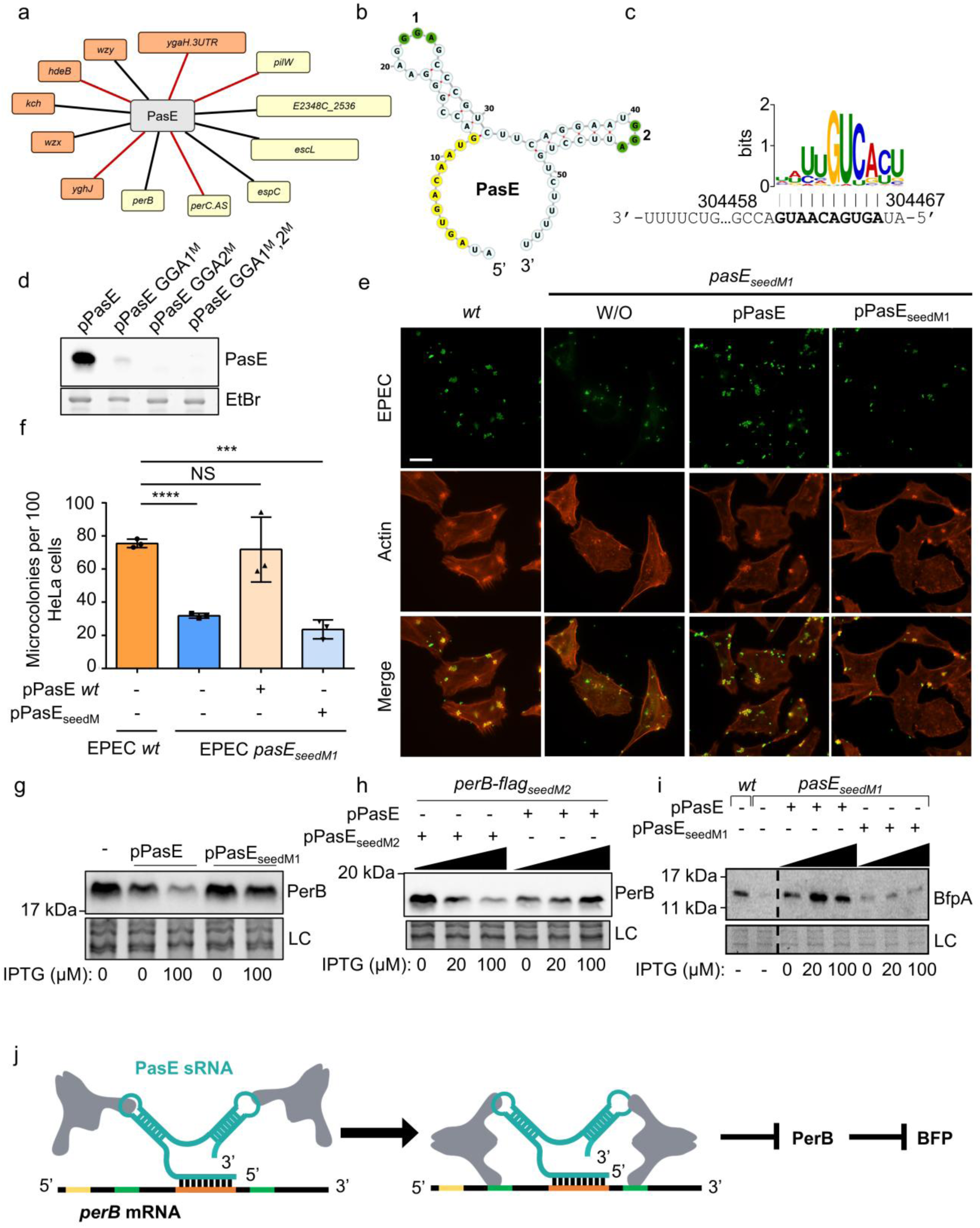
PasE is a CsrA-dependent novel sRNA repressing PerB production. (a) A segment of Figure 2 depicting the most stringent set of putative PasE target RNAs. (b) Predicted secondary structure of PasE. The seed sequence, involved in binding target RNAs, is marked in yellow and the two GGA motifs of the putative CsrA binding sites are marked in green and numbered 1 and 2. (c) Sequence logo of a motif shared by PasE putative targets aligned with its complementary PasE sequence, defined as its seed (bold fonts). The motif is presented in the 5’ to 3’ orientation. Vertical black and gray lines mark the potential base pairs, corresponding to strong and weaker motif positions, respectively. The genomic coordinates of the sRNA corresponding to the motif’s start and end are displayed. All of the 67 putative target RNAs, including the 12 shown in (A), contain the motif (E-value 4.7x10^-28^). (d) PasE stability is compromised upon perturbation of its CsrA binding sites. EPEC was transformed with plasmids expressing either wild-type PasE or PasE mutants under an IPTG-regulated promoter. Mutations were introduced in the predicted CsrA binding sites, including the first (GGA1^M^), second (GGA2^M^), or both sites (GGA1^M^, 2^M^) (Fig. S6a-b). Bacteria were grown to the exponential phase in DMEM, and the extracted RNA was analyzed by Northern blot. EtBr was used as a total RNA loading control. (e) Wild-type EPEC (*wt)*, or EPEC seed mutant (*pasE_seedM1_*) (strain LA9156, Fig. S6c), were used to infect HeLa cells in DMEM supplemented with IPTG. The *pasE_seedM1_* mutant remained untransformed (W/O), or complemented with a plasmid expressing wild-type *pasE* (pPasE), or a mutated *pasE* (pPasE_seedM1_). Infected cells were fixed after 135 minutes, stained for EPEC (green) and actin (red), and analyzed by fluorescent microscopy. Scale bar is 20 µm. (f) Quantification of the experiment shown in (e). The number of host-attached microcolonies in 100 randomly selected host cells was counted manually. Bars represent the mean of three biological replicates. The EPEC strains and supplemented plasmids used are indicated. Statistical significance was assessed using an unpaired t-test (*** p < 0.001; **** p < 0.0001; NS, not significant). (g-h) PasE-*perB* base pairing is required for the downregulation of *perB*. (g) EPEC expressing PerB-FLAG (strain LA9372) was transformed with a plasmid expressing PasE (pPasE), or PasE_seedM1_ mutant (pPasE_seedM1_), or remained plasmid-free (-). Bacteria were grown in DMEM to an OD of 0.2, and expression of PasE or PasE_seedM1_ was induced after the first hour of growth with 0.1 mM IPTG. PerB-FLAG levels were detected by western blot using anti-FLAG antibodies. In panels g-i, stain-free total protein served as Loading Control (LC). (h) EPEC expressing *perB-flag* containing a synonymous mutation that disrupts the PasE seed binding site (termed *perB-flag_seedM2_*) was transformed with a plasmid expressing either the compensatory PasE_seedM2_ (pPasE_seedM2_), or wild-type PasE (pPasE). Strains were grown to OD 0.2, and sRNA expression was induced during the last 2 hours of growth with 0.02 or 0.1 mM IPTG. Proteins were extracted and analyzed as described in (g). (i) PasE promotes BfpA production. Wild-type EPEC (*wt*), or EPEC *pasE_seedM1_* with or without ectopic expression of PasE *wt* or PasE*_seedM1_* mutant, were grown in DMEM with IPTG as indicated. At OD ∼0.2, proteins were extracted and analyzed by Western blot using anti-BfpA antibody. Two segments of the same blot are shown (separated by a black dashed line). (j) Model for *perB* repression through PasE-CsrA cooperation. The PasE-CsrA complex is guided to *perB* mRNA via PasE-*perB* base pairing. This enables CsrA to bind to sites flanking the base-paired sequence, resulting in the repression of PerB production and consequently enhancing BFP production. The *perB* sequence complementary to the PasE seed is shown in orange, and potential CsrA binding sites in green (Fig. S8). The translation initiation site is marked in yellow.

### PasE regulates EPEC infection

The predicted PasE targets include virulence-associated genes: *pilW*, encoding a protein involved in host cell attachment^18^; *escL*, an essential component of the T3SS^31^; and *espC*, an autotransporter toxin^32, 33^ (Fig. 3a). This raised the question of whether PasE directly contributes to EPEC infection. Since all the putative targets of PasE contained a common motif complementary to the PasE seed sequence (Fig. 3c), we mutated the seed by replacing GAC with CCA sequence in the EPEC chromosome (termed *pasE_seedM1_*). This mutation is predicted to disrupt potential pairing with target mRNAs while preserving PasE size, secondary structure, and ability to bind CsrA (Fig. S6c-e). We then compared the ability of wild-type EPEC and EPEC *pasE_seedM1_* mutant to infect HeLa cells (Fig. 3e-f). The mutant exhibited substantial reduction in size and number of host-attached bacterial microcolonies, a phenotype typically related to the BFP function^20^. We also noted a reduction in the formation of actin structures termed pedestal in the infected cells, indicating tempered T3SS activity^34^. Complementation with a plasmid expressing wild-type PasE restored efficient infectivity, demonstrating that PasE enhances EPEC virulence. To explore how PasE promotes virulence, we focused on its impact on the expression of *perB,* since *perB* is one of the strongest predicted PasE targets (Supplementary Table 2). Furthermore, we found that PerB antagonizes EPEC infectivity by repressing the expression of BFP (Fig. S7a-c), suggesting that PasE promotes EPEC infectivity by *perB* repression.

### PasE-CsrA cooperation is required for *perB* repression

To test the prediction that PasE represses *perB* through a base pairing mechanism, we overexpressed PasE or PasE_seedM1_ and observed that PasE, but not PasE_seedM1_, causes a decrease in PerB levels (Fig. 3g). Importantly, overexpression of PasE did not affect the levels of PerA, a positive autoregulator encoded on the same operon (Fig. S7d). Reciprocally, wild-type PasE failed to repress *perB* carrying a synonymous mutation that disrupted the complementary sequence (termed *perB_seedM2_*) (Fig. 3h). Introducing a compensatory mutation in PasE (termed PasE_seedM2_) restored repression of *perB_seedM2_* (Fig. 3h, Fig. S6c-d). These results suggest that PasE represses PerB production via base pairing. Consistent with this, expression of PasE, but not PasE_seedM1_, increased BFP expression (Fig. 3i). Overall, these results indicate that PasE represses PerB production through a mechanism involving direct base pairing (Fig. 3j).

The role of CsrA in PasE-mediated *perB* repression appears to be multifactorial. First, the stabilization of PasE by CsrA (Fig. 3d and Fig. S4f) is clearly critical for *perB* repression. In addition, *perB* contains several putative CsrA binding sites (Fig. S8b). Moreover, the RIL-seq data show that CsrA binds to one of these sites to form a ternary complex with PasE and *perB* mRNA (Supplementary Table 2). Yet, CsrA-*perB* singles (binary complexes) were not detected (Supplementary Table 3). These results indicate that CsrA binds to the *perB* mRNA in a PasE-dependent manner, possibly because these sites become available to CsrA only after base pairing with PasE (Fig. S8c-d), or due to the low affinity of CsrA to these sites. Taken together, our data suggest that base pairing between PasE and *perB* guides subsequent binding of the PasE-associated CsrA to the adjacent CsrA binding sites, thereby inhibiting *perB* expression.

### Modified PasE redirects CsrA to a designated target mRNA

To explore whether PasE can be engineered to guide CsrA to repress new targets, we employed as an initial test case the well-characterized CsrA target, *nleA*^13^. NleA repression involves the initial binding of one CsrA subunit to a high-affinity site on the *nleA* mRNA 5’UTR, which facilitates the consequent binding of the second subunit to a lower-affinity site overlapping the ribosome binding site, thereby inhibiting translation (Fig. 4a)^13^. Importantly, the initial binding to the high-affinity binding site is critical for repression, as in an EPEC mutant lacking this binding site (*5’utr_mut_-nleA*), NleA production is no longer repressed (Fig. 4b and 4d). To restore NleA repression in this mutant, we constructed a PasE variant (PasE_seedNleA_) that contains a seed sequence designed to base pair with the region just upstream of the *nleA* ribosome-binding site in the *5’utr_mut_-nleA* mutant. This allows the variant to “replace” the function of the original high-affinity CsrA binding site (Fig. 4c, Fig. S9a-c). Remarkably, expression of PasE_seedNleA_ restored *nleA* repression, while a PasE mutant, which cannot base pair with the *5’utr_mut_-nleA* (PasE_seedM1_), failed to do so (Fig. 4d). To test whether this repression requires direct base pairing, we deleted 10 nucleotides comprising the PasE seed target site in the *5’utr_mut_-nleA* and observed no repression (Fig. 4e-f). This indicates that the base pairing is essential for *nleA* repression by PasE_seedNleA_. To test whether this base pairing is sufficient for the repression, we overexpressed a synthetic sRNA, JV300, with a seed identical to that of PasE_seedNleA_ (JV300_seedNleA_), but lacking a CsrA binding site^35^. The overexpressed JV300_seedNleA_ failed to restore *nleA* repression (Fig. 4g-h, Fig. S9c-d), indicating that base pairing *per se* is insufficient for *nleA* repression by PasE_seedNleA_. Taken together, these results reinforce the premise that by base pairing with its target, PasE_seedNleA_ facilitates the binding of the PasE-associated CsrA to the low-affinity binding site that overlaps the *nleA* ribosome binding site, thereby repressing *nleA* expression (Fig. 4c, d).

**Figure 4.**
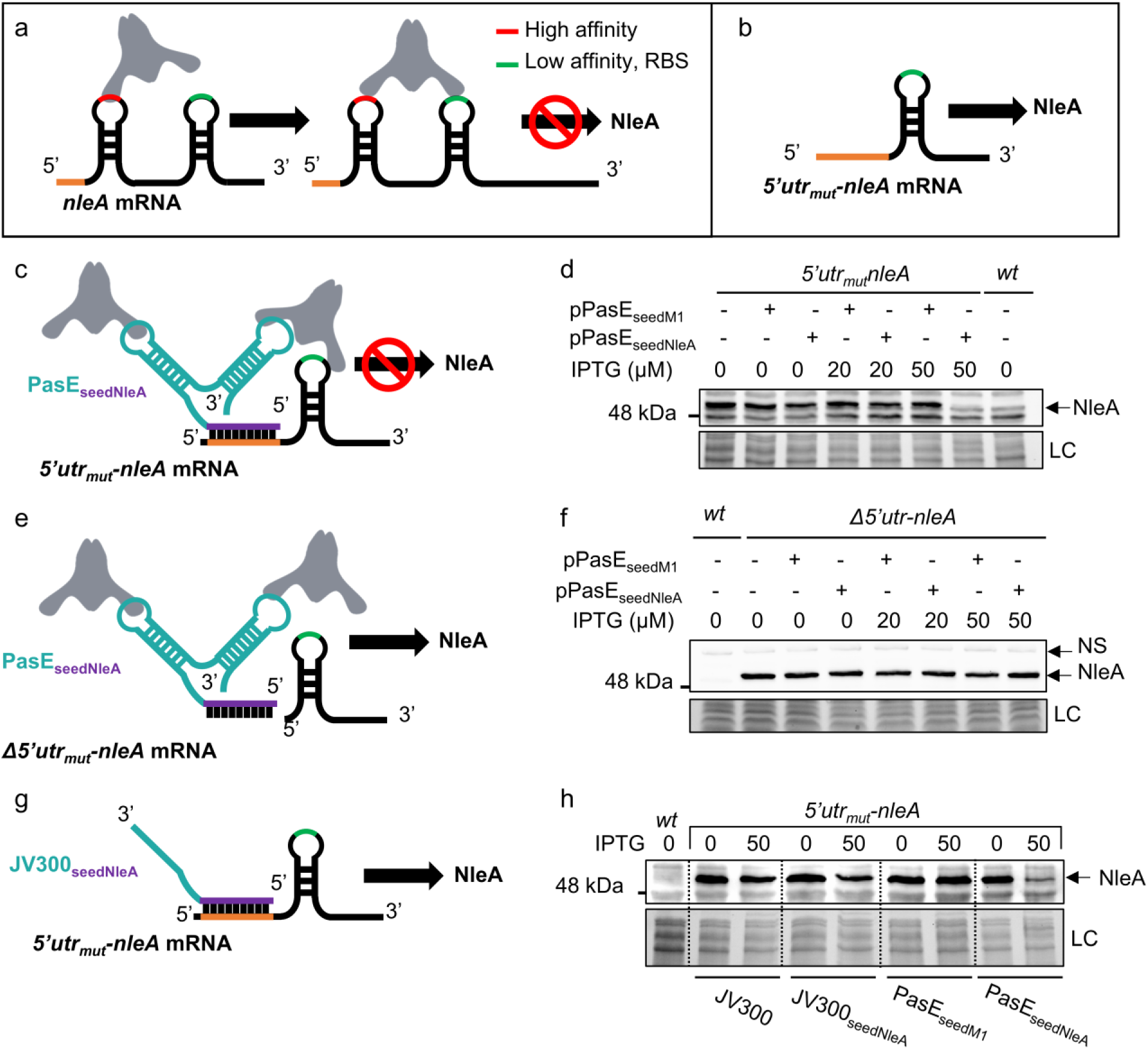
Modified PasE redirects CsrA to a designated target mRNA. (a) CsrA was reported to repress NleA via the canonical model. CsrA first binds to a high-affinity site (red) in the 5’UTR of *nleA*, facilitating the consequent interaction of its second subunit with a weak CsrA binding site that overlaps the ribosomal binding site (green), thereby repressing NleA production ^1, 13^. (b) Upon deletion of the high-affinity CsrA binding site in the *nleA* 5′ UTR (the mutant termed *5’utr_mut_-nleA*), CsrA can no longer repress NleA production ^13^. (c-d) Programmable mechanism of gene repression by CsrA-PasE cooperation. (c) Illustration of how modifying the PasE seed sequence re-targets CsrA to a new target mRNA (*nleA*), restoring repression even in the absence of a high-affinity CsrA binding site. PasE carrying a modified seed sequence (purple) base pairs with the 5’ *utr_mut_*-*nleA* mRNA (orange). This interaction enables the recruited CsrA to bind the low-affinity CsrA binding site overlapping the ribosomal binding site (green), thereby restoring gene repression. (d) NleA repression by programmed PasE, as illustrated in (c). Wild-type EPEC (*wt*) and *5’utr_mut_-nleA* mutant were transformed, or not, with plasmids expressing PasE variants whose seeds either complement the *5’utr_mut_-nleA* sequence (pPasE_seedNleA_), or do not (pPasE_seedM1_). Bacteria were grown in DMEM for 3 hours to OD 0.2, and when indicated, IPTG was added during the final two hours. Proteins were extracted and analyzed by Western blot using an anti-NleA antibody. The bands below and above the NleA band represent nonspecific cross-reactivity. LC, loading control, total protein staining. (e-f) Regulation by programmed PasE requires base pairing. (e) Schematic illustrating the loss of complementarity in the EPEC *Δ5′utr-nleA* mutant, which lacks the PasE_seedNleA_ seed-complementary sequence (see also Fig. S9c). (f) Wild-type EPEC (*wt*) and the Δ*5’utr-nleA* mutant were transformed, or not, with a plasmid expressing PasE_seedM1_ or PasE_seedNleA_, as indicated. Cultures were grown and analyzed as in (d). The NleA band is indicated by an arrow. NS-nonspecific band. LC-loading control. (g-h) Base pairing *per se* is insufficient for regulation by programmed PasE. (g) Schematic illustrating that a sRNA harboring a seed sequence but not CsrA binding sites does not affect *nleA* levels. (h) Wild-type EPEC (*wt*) or a *5’utr_mut_-nleA* mutant, each harboring plasmids expressing JV300, JV300_seedNleA_, PasE_seedM1_, or PasE_seedNleA_ (pJV300, pLA12635, pLA9194, and pLA11872, respectively) were grown in DMEM for 3 hours to an OD 0.2. sRNA expression was induced by adding IPTG (50 µM) after the first hour of growth. Proteins were extracted and subjected to Western blot analysis using anti-NleA antibody. LC, loading control, total protein.

### Repression of an antibiotic resistance gene by modified PasE

Typical ribosome binding sites contain a GGA sequence^28, 36^ and thus are potential low-affinity CsrA binding sites. We therefore predicted that various genes regularly not targeted by CsrA could be repressed by CsrA when guided by customized PasE variants. To test this prediction, we selected the *cat* gene, encoding chloramphenicol acetyltransferase, leading to chloramphenicol resistance. This gene is not known to be repressed by CsrA, and its ribosome binding site contains a GGA motif (Fig. 5a). We expressed in an EPEC strain carrying a chromosomal *cat* gene, five PasE variants (PasE_seedCAT1-CAT5_) whose seed regions are complementary to sequences located upstream of the *cat* ribosome binding site and the encompassed putative CsrA-recognition GGA motifs (Fig. 5b). Remarkably, three PasE variants markedly repressed *cat* expression, rendering the bacteria chloramphenicol-sensitive (Fig. 5c and Fig. S10). To determine whether *cat* repression requires both PasE-mRNA base pairing and CsrA binding, we generated an additional PasE variant (PasE_seedCAT6_) configured to impose either base pairing in the absence of CsrA binding, or CsrA association without base pairing (Fig. 5b). Notably, bacteria expressing this variant remained chloramphenicol-resistant, indicating that neither base pairing alone nor CsrA binding alone is sufficient for repression (Fig. 5c). We could not detect a correlation between *cat* repression and the seed melting temperature or the levels of the respective PasE in the bacteria (Fig. 5b and Fig. S9e-f). Yet, the repression ability correlated with an optimal distance between the GGA motif and the base pairing sequence on the *cat* 5’UTR (Fig. 5b-c and Fig. S10). These results suggest that PasE base pairing, positioned optimally relative to the GGA motif, may guide PasE-bound CsrA to engage this motif, thereby stabilizing the interaction of the PasE-CsrA complex with the *cat* mRNA. Collectively, our findings support the idea that PasE can be engineered to recruit CsrA to defined RNA sites, enabling context-dependent regulatory outcomes, including the re-sensitization of bacteria to antibiotic treatment.

**Figure 5.**
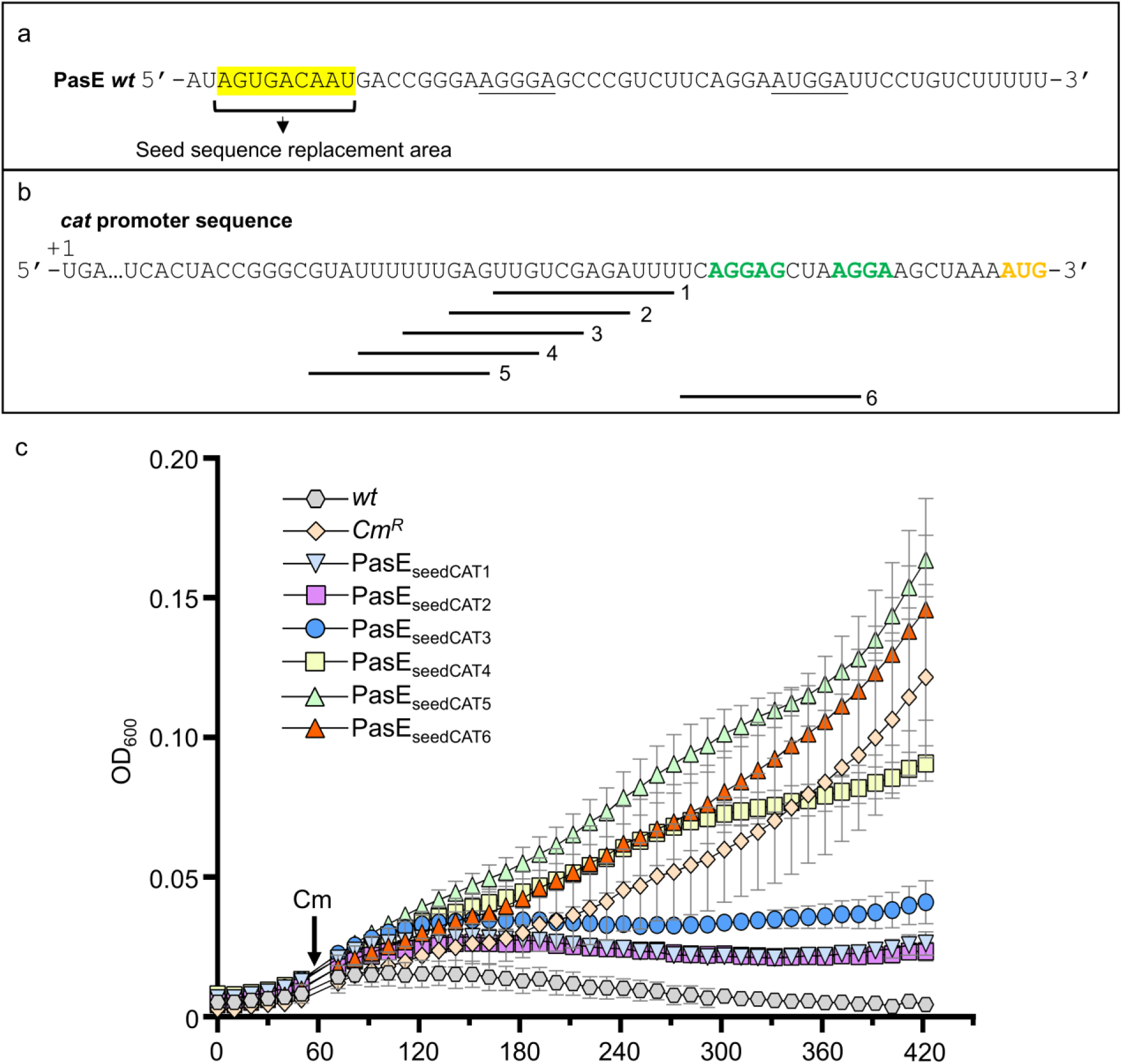
Repression of antibiotic resistance gene by programmed PasE. (a) The PasE sequence. The seed sequence is highlighted in yellow. (b) The 5’UTR of the *cat* gene ^42^. The +1 nucleotide is highlighted in yellow. GGA motifs corresponding to the ribosome binding site are shown in green, and the translation start site in orange. Lines numbered below the sequence mark the 12 nucleotide regions within the *cat* 5’UTR targeted for base pairing with PasE variants. For each region, the PasE seed sequence shown in (a) was replaced by a sequence complementing the corresponding *cat* 5’UTR sequence (Fig. S9f). These variants were designated PasE_seedCAT1_ to PasE_seedCAT6_. (c) EPEC containing a chromosomal *cat* gene was grown with or without plasmids expressing the modified PasE variants shown in (b). Bacteria were grown in LB at 37 °C in a 96-well plate within a plate reader, and OD was measured every 10 minutes. After 50 minutes of growth, chloramphenicol was added (50 µg/ml), as indicated. The used PasE variants are indicated below the plots. Wild-type EPEC was used as a chloramphenicol-sensitive control. The average of three biological replicates is shown. Error bars represent the Standard Error of the Mean (SEM). Student’s *t*-test between the final OD measurements of the chloramphenicol-resistant strain (*Cm^R^*) and the strains overexpressing PasE_seedCAT1-3_ was carried out, p-value < 0.0001 (****).

## Discussion

Here, we uncovered more than 800 CsrA-RNA ternary complexes involving CsrA bound to two distinct RNAs. Focusing on the PasE-CsrA-*perB* complex, we revealed an unexpected layer of post-transcriptional regulation in which the PasE sRNA guides CsrA to specific target mRNAs, thereby affecting EPEC virulence mechanisms. The regulatory effects observed for PasE-CsrA may likewise be extended to additional sRNA-CsrA-mRNA ternary complexes identified in our study (Fig. 2). Our findings provide new insights into the regulatory mechanisms underlying the actions of both sRNAs and CsrA. We show that, in addition to regulating their target mRNAs by direct base pairing, some sRNAs may regulate their targets by recruiting CsrA and guiding it to specific transcripts. Conversely, CsrA not only binds its targets directly but can also be guided by sRNAs to bind otherwise suboptimal sites on mRNAs. Together, these findings broaden our understanding of both CsrA- and sRNA-mediated regulation, uncovering regulatory layers that are likely widespread across bacteria, yet remain largely uncharted.

In many bacterial species, Hfq is considered to be the major mediator of post-transcriptional regulation, acting as an RNA chaperon that promotes sRNA-mRNA base pairing, thus leading to regulation of gene expression^37^. While we showed here a case where the sRNA guides CsrA to the target, it remains to be explored whether in other cases of ternary complexes CsrA functions similarly to Hfq, as a chaperon promoting sRNA-mRNA base pairing, which *per-se* accomplishes the regulation. Comparison of the RNA interactomes of CsrA and Hfq shows that these two global regulators engage sRNAs to form distinct, parallel post-transcriptional regulatory networks rather than a single, unified system (Fig. S11). Whether and how these networks are functionally coordinated merits further exploration, given that many sRNAs are bound by both CsrA and Hfq^23^. Our data also reveal the occurrence of ncRNA-CsrA-ncRNA complexes, some of which are highly abundant. The functions of most of the involved ncRNAs are yet unknown, although several are putative or previously characterized sRNAs. Whether this unique architecture represents an additional regulatory mechanism warrants further study. An additional notable observation is that most sRNAs displayed a pronounced bias toward localization at either the 3′ or 5′ end of chimeric fragments (Fig. S2). The biological significance of this bias remains unclear. By contrast, in Hfq RIL-seq experiments sRNAs were preferentially detected at the 3′ end of chimeras^26^, consistent with the structural organization of the sRNA–Hfq complex^38^. Cumulatively, our analysis opens up many new, novel, and exciting possible lines of investigation.

In addition to the ternary complexes, we identified more than 1,200 CsrA–RNA binary complexes. Interestingly, comparing the binary CsrA-RNA interactome of EPEC with that of the nonpathogenic *E. coli* K-12^2^, revealed a high degree of overlap. Over 88% (250) of CsrA targets were shared between the EPEC and K12 datasets, underscoring the robustness and conservation of the canonical CsrA-RNA interactions (Supplementary Table 3). Yet, we identified 450 additional binary complexes in EPEC, likely reflecting the EPEC larger genome, encoding many virulence genes^39^. Some of these genes were shown to be regulated via the canonical CsrA–RNA interactions^11, 13, 40, 41^ and our dataset now provides a framework for further exploration of CsrA-mediated regulation of virulence via this canonical mode of action.

Finally, our results also establish CsrA as a regulatory factor whose activity extends beyond direct mRNA binding and can be dynamically redirected by sRNAs. This sRNA-CsrA partnership can be rewired to control new targets, opening opportunities for programmable RNA circuits with potential applications in RNA therapeutics and antimicrobial resistance management.

## Declaration of generative AI and AI-assisted technologies in the manuscript preparation process

During the preparation of this work the authors used ChatGPT in order to verify the text grammar and find suitable synonyms. After using this tool/service, the authors reviewed and edited the content as needed and take full responsibility for the content of the published article.

## Acknowledgments

We thank Gad Frankel (Imperial College) and Samantha Gruenheid (McGill) for providing antibodies, Nathalie Q. Balaban (The Hebrew University of Jerusalem) for helpful advice, for HM, SBY and IR group members for helpful discussions and Roy Chaudhuri (Sheffield Bioinformatics Core) for help with gene annotations.

## Author contributions

Conceptualization: IR, HM, NE, LA, RFR, SBY

Methodology: RFR, LA, NE, HM, IR

Investigation: RFR, LA, NE, SF, YA, LA, MB, NK, YX

Visualization: LA, RFR

Funding acquisition: HM, IR, SBY

Administration MR

Supervision: HM, IR

Writing – original draft: RFR, LA, NE, YA, HM, IR

Writing – review & editing: RFR, LA, NE, SF, YA, LA, MB, NK, YX, SBY, HM, IR

## Data availability

The sequencing results of the RIL-seq libraries are available in ArrayExpress, with accession E-MTAB-13514.

## Methods

### Strains, plasmids, primers, and basic procedures

Bacterial strains, plasmids, primers/probes and antibodies are listed in Supplementary Information. Strains were constructed using the lambda red system, as previously described ^42, 43^. Briefly, EPEC cultures expressing the lambda red proteins were electroporated with a DNA fragment containing a selective antibiotic cassette or the *tetA-sacB* cassette followed by second round of transformation with the DNA fragment containing the desired genetic manipulation, all fragments contained 25-50 nucleotides flanking region of homology to the target gene. Next, the genetic manipulation was verified by PCR of the recombination target site followed by DNA sequencing.

Luria-Bertani (LB) medium was used for overnight growth and supplemented, when necessary, with antibiotics at the following concentrations: ampicillin (Amp, 100 μg/ml), streptomycin (Strep, 50 μg/ml), chloramphenicol (Cm, 25 μg/ml), tetracycline (Tet, 20 μg/ml), or kanamycin (Kn, 40 μg/ml). To mimic infection conditions, bacteria were statically grown overnight in LB at 37°C, then sub-cultured by diluting 1:50 with Dulbecco’s modified Eagle medium (DMEM, Biological Industries). All strains expressing PasE or its mutants also harbor a plasmid expressing LacI^q^ (pREP4, Qiagen) to repress PasE expression. Unless otherwise indicated, strains were grown for three or four hours in DMEM. When needed, IPTG (isopropyl-D-thiogalactopyranoside, Sigma) (0.1 mM) was added during the last two hours of growth.

### RIL-seq experiment

Single colonies of wild-type EPEC (E2348/69) and *csrA-3xflag* (NN6861) strains were inoculated from a fresh LB plate in liquid LB and grown over-night without shaking at 37°C. Bacteria were then diluted (1:50) in 200 mL of pre-warmed DMEM and grown to an OD_600_ 0.4. The bacteria were harvested and subjected to RIL-seq experiments as described ^25, 26^ with modifications to several steps of the protocols (step numbers according to Melamed et al. ^25^): Step 10: imidazole was not added to the wash buffer used throughout the protocol. Step 5: cells were exposed to 800mJ of UV irradiation. Step 22 for non-specific binding and Step 23 for binding between magnetic beads and antibody were not carried out since beads containing FLAG antibody were used (Anti-FLAG M2 Magnetic beads, M8823, Sigma). Step 25: 60 µl anti-FLAG beads were used for immunoprecipitation. Step 28: The first two washes were carried out with a wash buffer containing 1M NaCl and the remaining three washes were carried out with the wash buffer as indicated in Reagent Setup at Salt solution for wash buffer section ^25^, all washes were carried out at volume of 500 µl. Step 29: The RNase A/T1 mixture was diluted 1/40 in water. Step 31: The RNase digestion was carried out for five minutes. The experiments were performed in six biological replicates for each strain. Sequencing of libraries was carried out by the RNAtag-Seq method ^44^ with the modifications described in Melamed et al. ^25^. Reads were mapped to EPEC E2348/69 genome version 19 (chromosome NC_011601.1 and three plasmids NC_011602.1, NC_011603.1, and EU580135) with additional annotations defined in Pearl Mizrahi et al. ^23^. Computational analysis and identification of statistically significant chimeras (S-chimeras) were carried out as described in Melamed et al. ^25^.

### Computational analysis for the identification of statistically significant single RNA molecules bound to CsrA

To identify single RNA molecules bound on CsrA, we ran the RIL-seq code: significant_regions.py (https://github.com/asafpr/RILseq) with the two parameters: -- all_interactions, and --only_singles, for each replicate library for both wild-type and CsrA-FLAG libraries. Counts of reads mapped to overlapping sequences of the same annotation were summed. DESeq2 ^45^ was carried out for comparing corresponding single reads of the six CsrA-FLAG libraries and the six wild-type libraries. Entries with a base mean greater than five, p-adjusted value lower than 0.05, and log2FoldChange value greater than 2 were considered as RNAs that are statistically significantly enriched on CsrA.

### Calculation of the number of MNGGA motifs in single RNA molecules (Supplementary Table 2 and 3)

RNA sequences were extracted based on the genome coordinates of the chimeric fragments with an expansion of 50 nucleotides upstream and downstream. We looked for the regular expression MNGGA (M stands for A/C and N stands for any nucleotide) ^2^ in each sequence and counted the number of appearances of this motif. To calculate the average probability of positional accessibility of the motif in the folded RNA, each MNGGA motif sequence was expanded by nine nucleotides upstream and downstream. The probability of accessibility was computed using RNAfold ^46^, and indicated in Supplementary Table 2 and 3.

### Generating CsrA RNA-RNA regulatory network

Based on our previous findings ^29^, we considered a consistently high abundance of reads corresponding to a specific sRNA-mRNA chimera across multiple libraries as a strong predictor of an interaction that leads to a regulatory outcome. To evaluate the repeatability, we regarded the number of libraries (*i.e*., biological replicates) in which each chimera was detected. We included in the CsrA RNA-RNA network RNA pairs identified in at least four libraries, and presented it using Cytoscape ^47^. Genes were classified as described ^39^.

### Pulldown of CsrA-SBP

EPEC *hfq-flag* (SPM6777) transformed, or not transformed, with a CsrA-SBP (Streptavidin Binding Peptide) expressing plasmid (pNN6431), were grown overnight at 37 ^°^C in LB or LB supplemented with Amp. The next day, bacteria were diluted 1:100 into 50 ml of pre-warmed DMEM and grown for 1h at 37^°^C, then IPTG (0.1mM) was added to induce CsrA-SBP expression and bacteria were allowed to further grow to an OD_600_ ∼0.5. Bacteria were then processed according to the RIL-seq methodology as described above. Briefly, the CsrA-SBP induced culture was centrifuged, washed trice in ice-cold PBS (Biological Industries 02-023-5A) and subjected to UV crosslinking, applying 800 mJ to crosslink CsrA-RNA molecules. The cross-linked cells were loaded onto Zirconium oxide beads 0.15 mm (Ornat Cat.ADV-ZrOB015), lysed (Retch, model no. MM400, tube adaptors, cat. no. 22.008.0008) and the cleared extracts were loaded onto 25 ul of anti-SBP beads (Cat-S1638-1ML Sigma Aldrich) for 90 minutes at 4 ^°^C. Beads were then washed twice with 1 M NaCl in washing solution (50mM sodium phosphate, 0.1% IGEPAL, Protease inhibitor cocktail, RNase inhibitor 0.1 U/μl) and three times with 300 mM NaCl in washing solution. The washed beads were then treated with RNase solution for 5 minutes at room temperature, washed, and subjected to Superase treatment. The treated beads were mixed with Laemmli sample buffer (Bio-rad cat 1610747), treated for 10 minutes at 95 ^°^C, and the denatured proteins were loaded into a 12% free-stain gel (Bio-rad Cat. 4568046) for further analysis by western blotting with anti-Flag and anti-SBP antibodies.

### Common motif analysis in the target set of each sRNA

The MEME suite ^30^ was used for the identification of a common motif in each set of bound RNAs associated with a known or putative sRNA. Genomic coordinates of the bound sequences were derived from the positions of the corresponding mapped reads in the RIL-seq data, extended by 20 nucleotides upstream and downstream of the RIL-seq original coordinates. MAST was then used to identify the complementary site of the motif on the relevant sRNA^48^.

### RNA secondary structure prediction

RNA secondary structures were predicted using Vienna RNAfold ^46^ and illustrated using Forna ^49^.

### Rapid amplification of cDNA ends (RACE) experiments

50 µg of total RNA were DNase-treated using the TURBO DNase kit (Invitrogen, cat. No. AM2238) in a total reaction volume of 100 µl, according to the manufacturer’s instructions. RNA was precipitated in ethanol and resuspended in 40 µl of RNase-free water. For 5’ RACE experiments, 15 µg of DNase-treated RNA were subjected to a treatment of RNA 5’ Pyrophosphohydrolase (RppH) (New England Biolabs, cat. No. M0356S), using 75 units of RppH in 1X reaction buffer, in a total reaction volume of 150 µl. The reaction was carried out for 1 h at 37°C. In parallel, 15 µg of DNase-treated RNA were subjected to a control reaction, incubated under the same conditions but without RppH. The RNA was precipitated in ethanol with sodium acetate and resuspended in RNase-free water. The RNA (both RppH-treated and non-treated) was incubated with 500 pmol of a 5’ adapter (/5InvddT/rGrUrG rArCrU rGrGrA rGrUrU rCrArG rArCrG rUrGrU rGrCrU rCrUrU rCrCrG rArUrC/, IDT) at 75°C for 2 min and then placed on ice. Ligation of the adapter to the RNA was carried out using 65 units of T4 RNA ligase 1 (New England Biolabs, cat. No. M0437M) in a total reaction volume of 25 µl containing 1X reaction buffer, 9% DMSO, 1 mM ATP, 20% PEG 8000 and 30 units of recombinant RNase Inhibitor (RRI) (Takara, Cat. No. 2313A), incubated at 22°C for 2h. Ligation products were treated with phenol and chloroform, precipitated in ethanol, resuspended in RNase-free water, and incubated with 50 µg of random hexamers for 5 min at 65°C, followed by fast cooling on ice. For first strand cDNA synthesis, the SuperScript III kit (Invitrogen, cat. No. 18080-044) was used according to the manufacturer’s instructions. cDNA was PCR amplified using primers 5’ RACE F and 5’ RACE R (CTC CCT TCC CGG TCA TTG TCA C and GTG ACT GGA GTT CAG ACG TGT GCT CTT CCG ATC, respectively (IDT)) and 2X PCRBIO HS Taq mix Red solution. For 3’ RACE, 15 μg of DNase-treated RNA were treated with Fast AP (Thermo Scientific, cat. No. EF0651) for 15 min at 37°C. The RNA was then treated with phenol and chloroform, ethanol precipitated, and resuspended in 5 μL of RNase-free water. The RNA was ligated to a 3’ adapter (5rApp/AGA TCG GAA GAG CGT CGT GTA GGG AAA GA/3ddC/, IDT) using 520 units of T4 RNA ligase 2 truncated K227Q (New England Biolabs, Cat. No. M0351) in a total reaction volume of 15 ul containing 1X ligase buffer, 20% PEG 8000 and 72 units of RRI for 2 h at 22°C. Ligation products were treated with phenol and chloroform, precipitated in ethanol and then incubated with 3’ RACE RT primer (ACA CTC TTT CCC TAC ACG ACG CTC TTC CGA TCT, IDT) 5 min at 65°C, followed by fast cooling on ice. For first strand cDNA synthesis, the SuperScript III kit was used according to the manufacturer instructions. cDNA was PCR amplified using primers 3’ RACE F and 3’ RACE R (GTG ACA ATG ACC GGG AAG GGA G and ACA CTC TTT CCC TAC ACG ACG CTC TTC CGA TCT, respectively (IDT)) and 2X PCRBIO HS Taq mix Red solution. The 3’ RACE and 5’ RACE amplified cDNAs were run on 3% Nusieve gel, and fragments of about 75 bp and 100 bp, respectively, were extracted from gel, cloned into pDrive cloning vector using the PCR cloning kit (QIAGEN) and transformed into competent Stellar cells (Takara, cat. No. 636763), which were plated on ampicillin LB plates. Ampicillin resistant colonies were subjected to colony PCR with M13 F and M13 R primers (GTTTTCCCAGTCACGACG and CAGGAAACAGCTATGAC, respectively), and the PCR products were Sanger sequenced.

### Northern blot

Total RNA samples (30 µg) were denatured for 10 min at 65 °C in loading buffer containing 65% formamide, separated on 7 M urea/6% polyacrylamide gels in 44.5 mM Tris-base, 44.5 mM Boric acid and 2 mM EDTA pH 8.0, and transferred to Zeta-Probe membrane by electroblotting. To detect wild-type PasE and seed mutants (PasE_seedmut1_ and PasE_seedmut2_) the membrane was hybridized with PasE specific [^32^P] or biotin 5’ end-labelled DNA probe (5’-ACAGGAATCCATTCCTGAAGACGGGCTCCCTTCCCGGT-3’). To detect PasE CsrA binding site mutants, a different biotin end-labelled DNA probe was used (5’-CTTCCCGGTCATTGTCACTAT-3’). In Fig. S4d, bacteria were grown to OD ∼0.6 (exponential phase) or OD ∼1 (stationary phase). In Fig. S4f, PasE expression was detected by growing the samples for six hours, after the first hour, 0.1 mM of IPTG was added to induce CsrA expression in the relevant sample. In Fig. 3d and Fig. S6d, bacteria were grown to OD ∼0.6 and PasE expression was induced after the first hour of growth.

### Microscopy and infection

HeLa cells were seeded in a 24-well plate (Nunc) at a density of ∼8x10^4^ cells per well and grown overnight in DMEM supplemented with 10% fetal bovine serum (FBS; Sigma) and antibiotics (penicillin-streptomycin solutions; Gibco). Next, the HeLa cells were infected with overnight statically grown bacteria at 37°C (MOI ∼1:100), followed by two hours and 15 minutes of infection at 37 °C in 5% CO2. When needed, infecting strains were supplemented with 0.02 mM IPTG as indicated. To terminate the infection, the cells were fixed with freshly prepared 3.7% formaldehyde, washed with PBS, perforated (0.25% Triton X-100 in PBS solution) for 20 min, washed again with PBS and stained for anti-EPEC (followed by secondary anti-rabbit Alexa 488; Cell signaling) and phalloidin-rhodamine (Sigma), and analyzed with fluorescence microscopy. Quantification of microcolony formation was performed using an unpaired t-test on experimental data based on three independent biological replicates, using GraphPad Prism version 6.07 for Windows (GraphPad Software).

### Western blot

Single colonies were grown from a fresh LB plate in liquid LB overnight without shaking at 37°C. Bacteria were diluted 1:50 from the overnight culture and grown in pre-warmed DMEM (01-053-1A, Sartorius) to exponential phase. Cells were pelleted and resuspended in Laemmli buffer (CAT. #1610747 Bio-Rad) with 100 mM DTT, boiled at 95 °C, centrifuged for 1 minute at 13,400 rpm at room temperature, and subjected to SDS-PAGE (Bio-Rad 12% TGX pre-cast gels, CAT #4561046). Loading amounts were normalized based on OD measurements of the cultures and further verified by using stain-free gel imaging (Bio-Rad). Antibodies are listed in Supplementary Information. In Figures 3g-i, S8A and S8D cells were grown to OD ∼0.2-0.3 and IPTG was induced after the first hour of growth. In Figures 4d, 4f and 4h, strains were grown to OD ∼0.4, and increasing concentrations of IPTG (0-50 µM) were added during the last two hours of growth.

### Conservation of *pasE*

The conservation analysis of the *pasE* sequence was done with BLASTn tool ^50^ available at https://blast.ncbi.nlm.nih.gov. As the *pasE* sequence is short, BLAST automatically adjusted the search parameters for short input sequences. Default BLASTn parameters were used, except for the following modifications: Max target sequences: 5000, Filtering and masking: disabled, Optimization: “Somewhat similar sequences (blastn)”, Database: “All genomes / Complete genomes”, Entrez query: bacteria[Organism] AND “complete genome”[Title]

The majority of high-scoring hits (96%) originated from *Escherichia coli* complete genomes (Supplementary Table 4). We note that the database may contain multiple genome variants of the same strain, e.g., *E. coli* K-12 MG1655 with different genome submissions (e.g., NZ_CP010443.1, NZ_CP010444.1). Alignments were processed from the BLAST output using the “Query-anchored with dots for identities” alignment output view. The *pasE* query sequence was placed on top, and all non-redundant hits were aligned against it. Identical hits were collapsed into a single representative, with the number of collapsed sequences indicated (Fig. S5).

### Chloramphenicol sensitivity assay

Culture was grown overnight without shaking at 37 °C, diluted 1:100 in LB into a 96-well plate (NunC delta surface, 150 µl/well), and grown in plate reader (SPARK, Tecan) set at 37 °C. OD_600_ was measured every 10 minutes. Chloramphenicol (Cm) was added after the first 50 minutes of growth. Wild-type EPEC was used as a Cm-sensitive strain, and EPEC harboring a *cat* gene (Strain NN6490) was used as a resistant strain. Cm sensitivity was assessed when expressing PasE variants that base paired different areas of the 5’ UTR of the *cat* gene.

### Disk diffusion assay

EPEC cultures were grown overnight at 37 °C without shaking and ∼5 x 10^6^ were plated (LB agar). Disks containing serial dilutions of chloramphenicol (Cm; 2 µg, 4 µg, 8 µg, 16 µg, 32 µg and 64 µg) were placed on the plates, which were then incubated overnight at 37 °C. Chloramphenicol sensitivity was assessed the following day. Wild-type EPEC served as the chloramphenicol-sensitive control.

## Supplemental information

**Supplemental information Figure 1.**
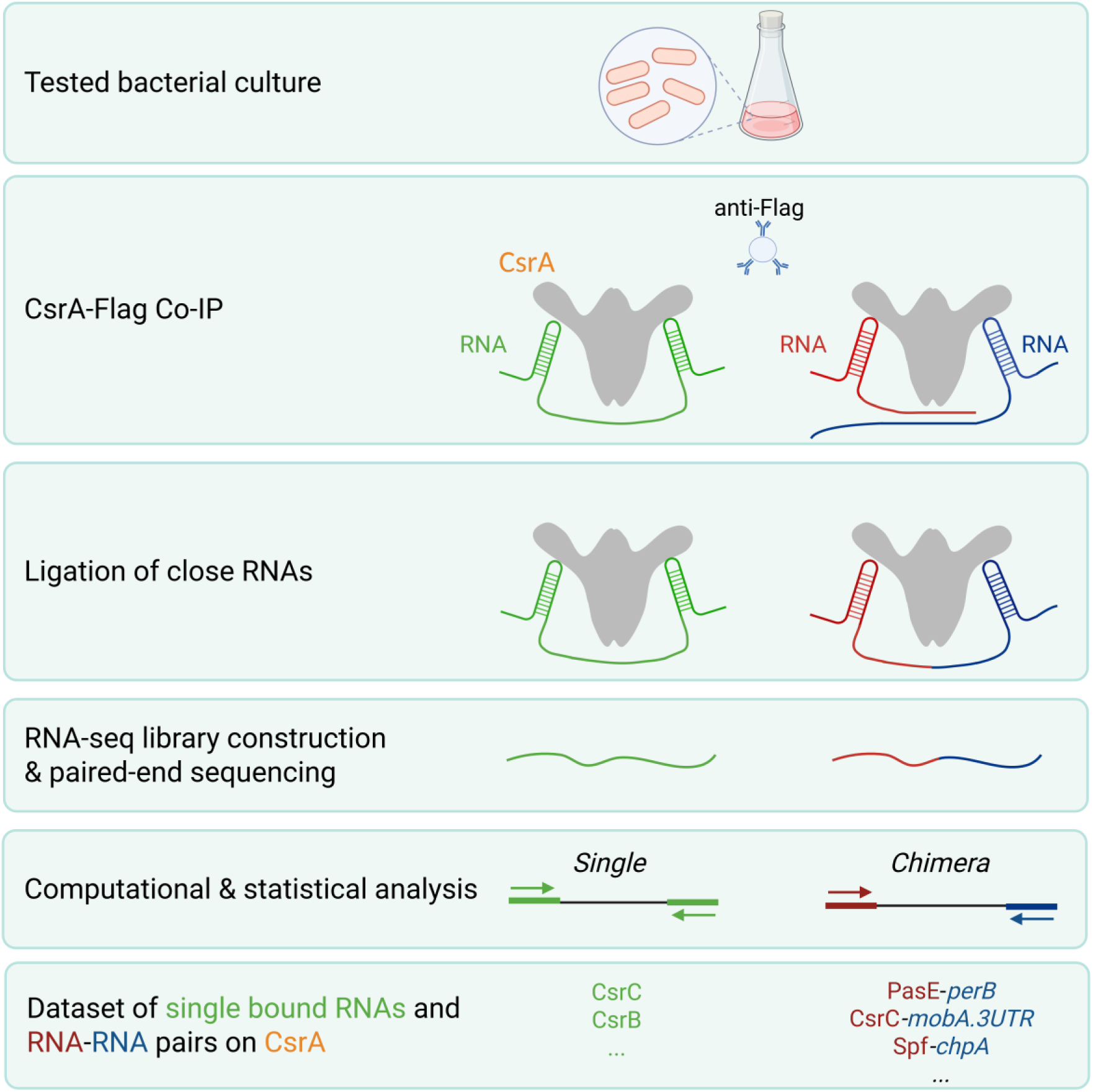
Main steps of RIL-seq applied to CsrA. EPEC expressing Flag-tagged CsrA was grown in DMEM to exponential phase and subjected to co-immunoprecipitation (co-IP) to capture CsrA together with its bound RNAs. RNAs bound to a CsrA dimer were pulled down (either single RNAs (green) or two distinct RNAs (red and blue)). The RIL-seq protocol ^25, 26^ included an RNA ligation step to ligate proximal RNAs, followed by digestion of the CsrA protein, RNA-seq library preparation, and paired-end sequencing. Computational analysis was then applied to identify single-fragment reads that map to one genomic location and chimeric reads that map to two distinct genomic locations.

**Supplemental information Figure 2.**
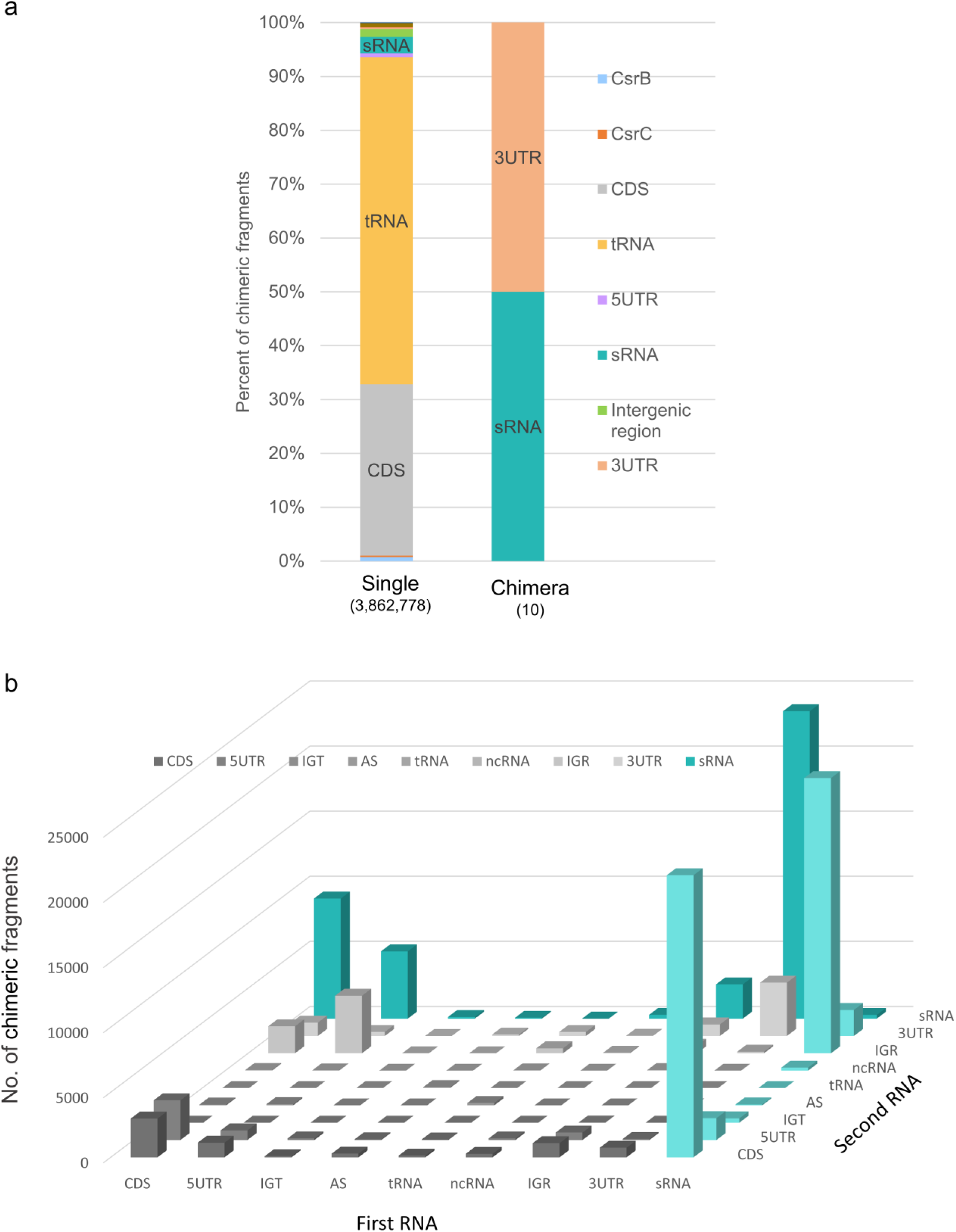
Background binding, comparison to CLIP-seq, and classification of CsrA-associated chimeric RNAs. (a) Distribution of single RNAs (left) and chimeric RNA fragments (right), nonspecifically absorbed on the magnetic beads when using wild-type EPEC (where CsrA is not FLAG tagged) in the RIL-seq experiment. The bound RNAs were categorized as in Fig. 1b to 5UTR, CDS, 3UTR, tRNA, ncRNA, AS (antisense), intergenic region (IGR), intergenic within transcript (IGT), CsrB, and CsrC. rRNAs were excluded from the analysis. The color code and fraction of each category (percent) are indicated. Total counts for detected single and chimeric RNAs are denoted in parentheses. Importantly, note that the total counts in this control (background binding) are very low compared to the RIL-seq results shown in Fig. 1b. (b) Distribution of the RNA pairs in chimeric fragments. The RNA fragments found in CsrA-associated chimeras were classified as in Fig. 1b (CDS, 5UTR, IGR, 3UTR, and sRNA). CsrC is included in the sRNA category, and CsrB is not included since it was not present in chimeras. The relative location of the fragments within the chimeras is indicated (First RNA=5’ and Second RNA=3’). The number of chimeric fragments for each category is indicated.

**Supplemental information Figure 3.**
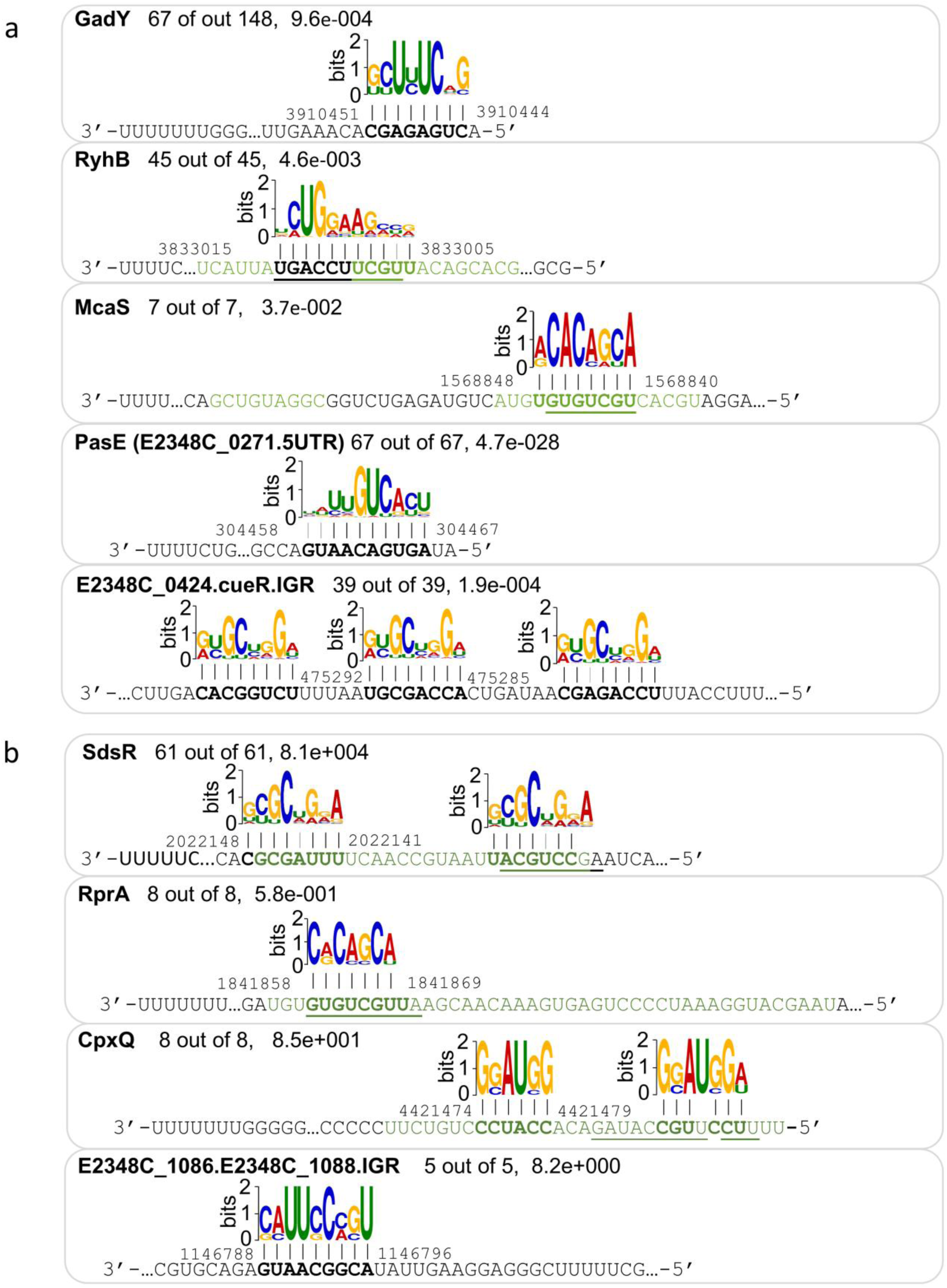
RNAs co-bound on CsrA with a specific sRNA share sequence motifs complementary to the respective sRNA. For each sRNA, the MEME-identified common motif in its target mRNAs is shown ^30^ in the 5’ to 3’ direction, aligned with its complementary site on the indicated sRNA, presented in the 3’ to 5’ direction. The number of sequences containing the motif (out of the total number of putative sRNA-bound sequences) and the MEME E-value ^48^, are indicated. Motif-sRNA base pairing is represented by vertical black or gray lines for strong or weak interactions, respectively. Complementary nucleotides in the sRNA are highlighted in bold. Previously reported Hfq binding regions ^26^ are marked in green, and sRNA nucleotides involved in base pairing supported by Hfq RIL-seq data are underlined. Genomic coordinates (EPEC E2348/69 genome) of the leftmost and rightmost base-pair positions on the sRNA are shown. The presence of shared motifs complementary to sRNA binding sites suggests base pairing between the sRNA and mRNA, supporting the reliability of our data. (a) RNAs co-bound on CsrA containing a statistically significant motif complementary to the respective sRNA. (b) RNAs co-bound on CsrA with a specific sRNA sharing a common motif complementary to the respective sRNA, but the motif is not statistically significant using the same threshold as in (a). For CpxQ, the binding site shown here corresponds to the one previously reported in ^51^.

**Supplemental information Figure 4.**
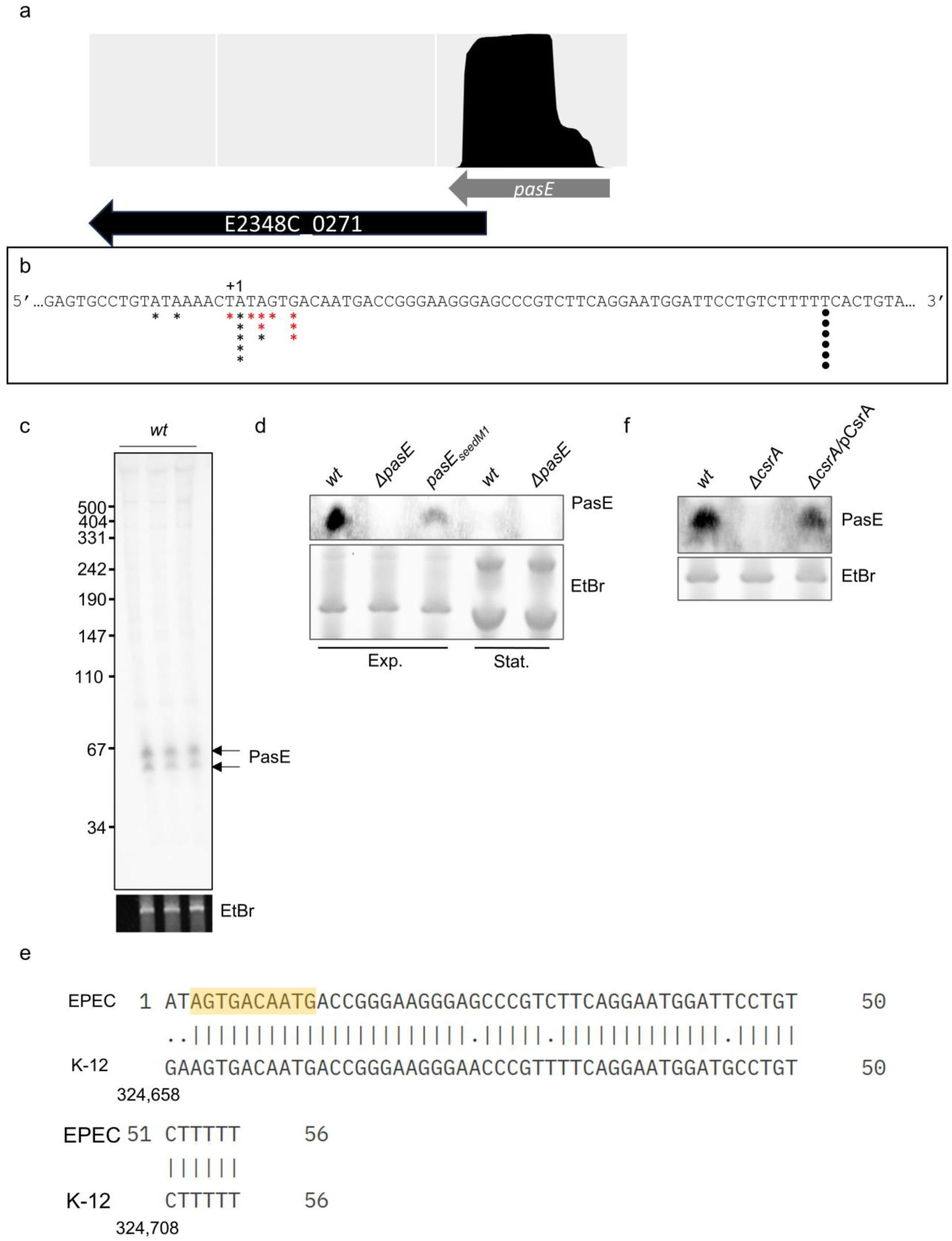
Characterization of the newly identified sRNA PasE. (a) Read coverage of PasE from an RNA-seq experiment of EPEC grown under infection-mimicking conditions ^23^. (b) Mapping the PasE 5’ and 3’ by RACE. Total RNA was treated, or not, with RppH, which converts tri-phosphate 5’ RNA ends into mono-phosphate 5’ ends. RACE experiments were done as described in Methods. Black and red asterisks indicate the 5’ positions of cloned and sequenced 5’ RACE products of RppH-treated and non-treated RNA, respectively. Black circles indicate the 3’ positions of cloned and sequenced 3’ RACE products. (c) PasE is stably expressed under host-mimicking conditions. Three biological repeats of EPEC *wt* grown to the exponential phase, RNA was extracted, and a Northern blot for *pasE* was carried out, using a radioactive probe. (d) Expression of *pasE* and *pasE_seedM1_* under host-mimicking conditions. Wild-type EPEC (*wt*), Δ*pasE* and *pasE_seedM1_* mutants were grown to the exponential growth phase (Exp.) in DMEM, RNA was extracted, and analyzed by Northern blot to detect PasE expression using a biotinylated probe. Wild-type and Δ*pasE* strains were also analyzed for PasE expression during the stationary phase (Stat.). (e) Alignment of PasE and the 3’UTR sequence of *ykgH* gene in *E. coli* K-12 MG1655. The seed sequence is highlighted in yellow. (f) The steady-state level of PasE is CsrA-dependent. Wild-type EPEC (*wt)* and a Δ*csrA* mutant with or without a plasmid expressing CsrA (pCsrA), were grown to the exponential phase under host-mimicking conditions. After the first hour of growth, ectopic expression of CsrA was induced for five hours. In panels c, d, and f, EtBr was used as a loading control for total RNA.

**Supplemental information Figure 5.**
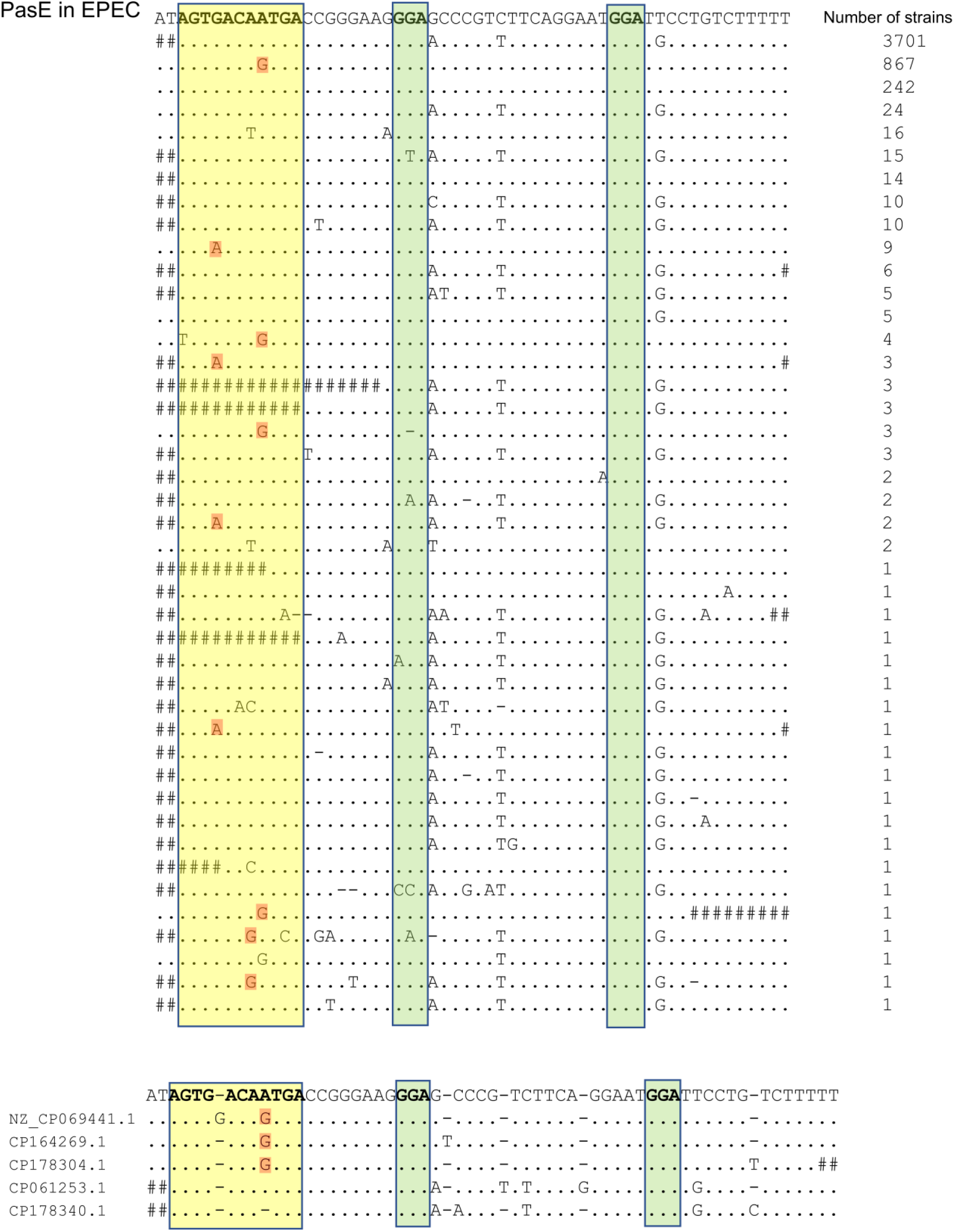
Conservation of the *pasE* sequence across ∼5,000 complete genomes of *E. coli* and *Shigella* strains. BLASTn alignments of the *pasE* sequence to *E. coli* and *Shigella flexneri* genomes are shown using the “Query-anchored with dots for identities” format. The EPEC *pasE* query is shown on top. Dots (.) represent identity to the query; mismatches are indicated by letters. Hits shorter than the query are padded with “#” at the edges. Identical sequences were collapsed and represented as one sequence. The number of collapsed sequences is indicated at the right of each alignment. Top panel: hits with no insertions. Bottom panel: five hits that contain insertions relative to the query, with corresponding GenBank accession numbers shown on the left. Sequences are shown as DNA (with thymine, T), as this is a genomic search. The *pasE* binding seed is highlighted in yellow and the two GGA motifs are shown in green. Orange rectangles indicate compensatory mismatches (e.g., G replacing A, maintaining potential RNA base pairing with U). Note that 22 genomes contained two hits for the *pasE* sequence and one genome contained three hits; therefore, the total number of genomes differs from the total number in Supplementary Table 4.

**Supplemental information Figure 6.**
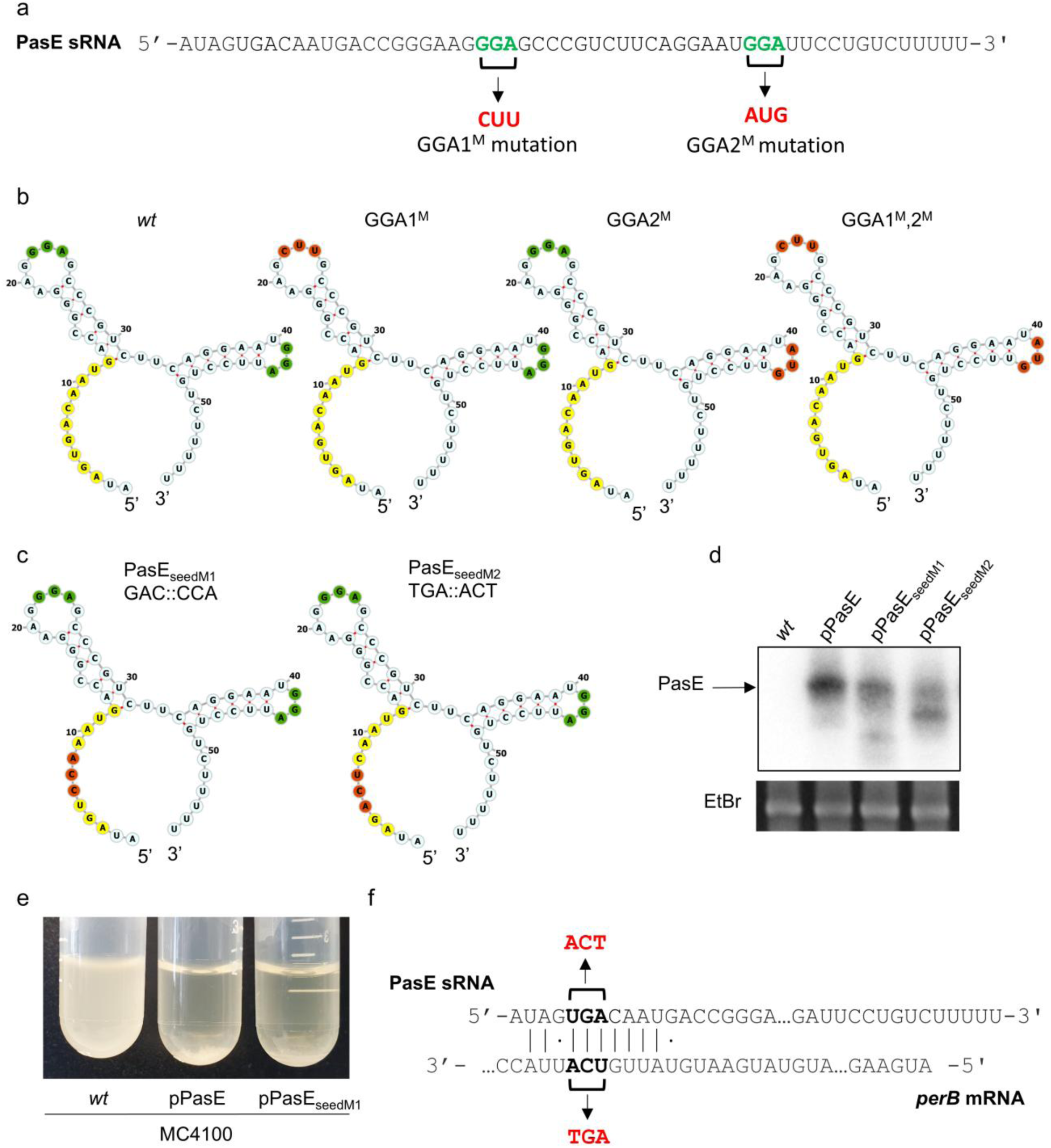
CsrA binding site and seed sequence mutations in PasE controls. (a) Location of the predicted CsrA binding sites of PasE, and their respective mutation. CsrA binding sites (GGA sequences) were marked in bold green, and the black arrow points to the respective mutation, marked in red. (b) Predicted secondary structure of the native and mutated PasE RNAs using RNAfold ^46^ and Forna ^49^. Native and mutant GGA sequences as well as the PasE seed are highlighted in green, red and yellow, respectively. The predicted mutated structures are similar to that of the wild type. (c) The secondary structures of the PasE seed mutations (seedM1 and seedM2) were predicted as in (b). The seed sequence is marked in yellow and the mutated nucleotides within it in red. The CsrA binding sites (green) are also highlighted. The predicted secondary structures of both PasE seed mutants remain similar to that of the native PasE. (d) Expression of recombinant PasE and PasE seed mutants. EPEC *perB-flag* was complemented with plasmids expressing wild-type PasE (pPasE), pPasE_seedM1_, or pPasE_seedM2_, or remained without the expression vector (*wt*). Bacteria were grown in DMEM for four hours to OD ∼0.2. PasE expression was induced by adding IPTG after the first hour of growth. EtBr staining of the total RNA was used as a loading control. A black arrow indicates the full-length transcript of PasE. The expression of the native chromosomally encoded PasE is not seen here (left lane) since the overexpression of the recombinant PasE variants dictate short exposure of the blot. (e) Overexpression of recombinant PasE or PasE_seedM1_, induces bacteria precipitation, typical to that caused by *pgaABCD* de-repression upon CsrA sequestering ^52^. *E. coli* K12 MC4100 (*wt*) or MC4100 expressing PasE (pPasE), or PasE_seedM1_ (pPasE_seedM1_) were grown overnight in LB within the tubes shown. Both PasE variants induced bacterial precipitation, likely via CsrA sequestering leading to *pgaABCD* expression. (f) PasE-*perB* predicted base pairing and compensatory mutation design for seedM2.

**Supplemental information Figure 7.**
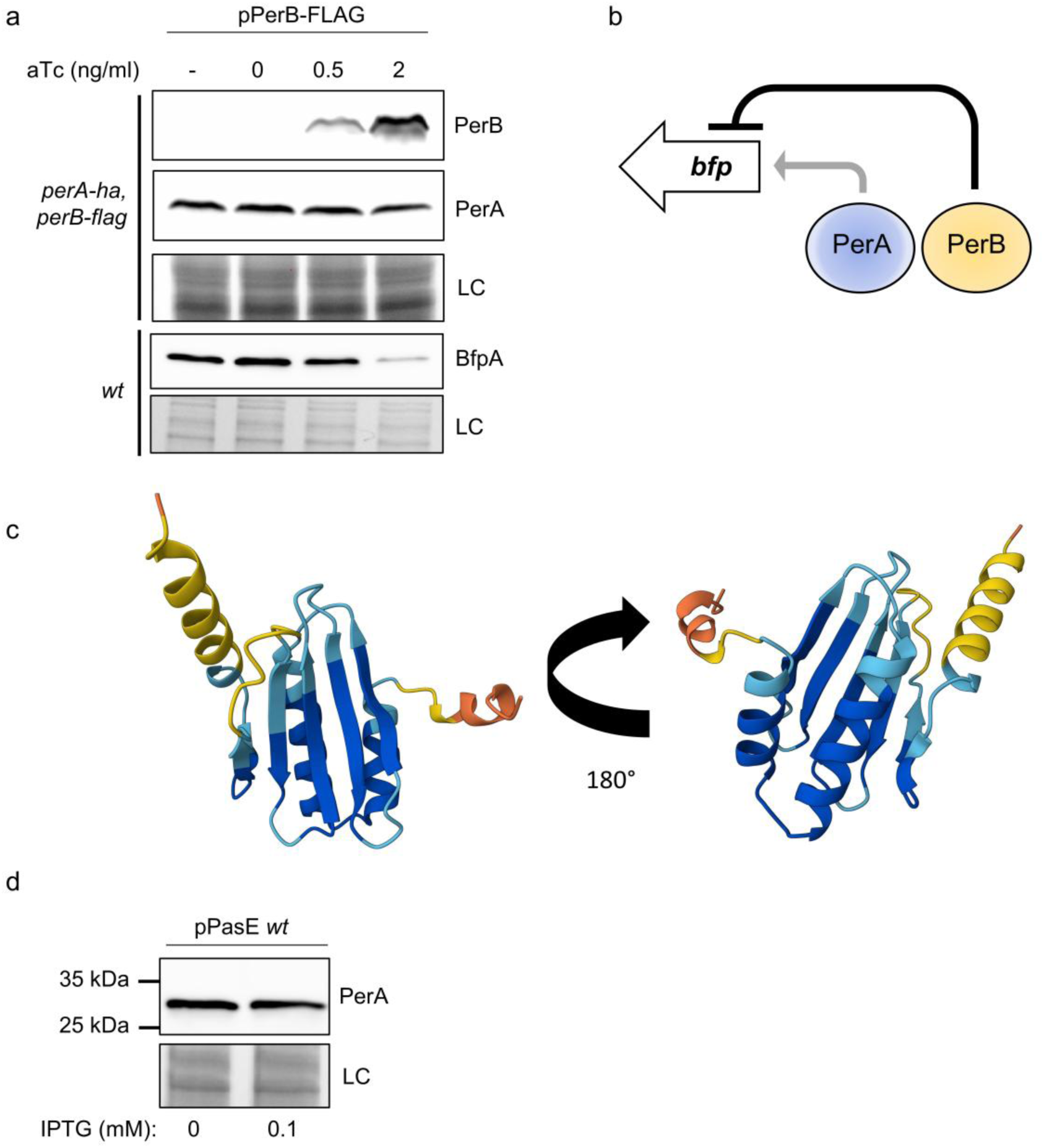
PerB represses BfpA but not PerA, and structural analysis supports a DNA-binding role for PerB. (a-b) PerB represses BfpA, but not PerA production. (A) EPEC *wt*, or EPEC *perA-ha*, *perB-flag,* both harboring a plasmid expressing PerB-FLAG were grown in DMEM supplemented with increasing concentration (0-2 ng/ml) of anhydrous tetracycline (aTc) and grown to the exponential growth phase. Western blot was performed using anti-HA, anti-FLAG and anti-BfpA for detection of PerA, PerB and BfpA, respectively. (b) An illustration of the negative regulation of BFP by PerB. Gray arrow indicates a previously characterized function of PerA’s role as a positive autoregulatory transcription factor ^53^. (c) Predicted structure of PerB. The AlphaFold model ^54^ of PerB shows with high confidence a beta-sheet core and helix-turn-helix, which are common features in many DNA-binding proteins. Computed Structure Models provide a per-residue confidence score (pLDDT) between 0 and 100. Some regions below 50 pLDDT may be unstructured in isolation. Structure colors are based on model confidence (very high; blue-pLDDT > 90, high; cyan-pLDDT > 70, low; yellow 50 < pLDDT ≤ 70, very low; orange ≤ 50). Image was taken from the Protein Data Bank (PDB) website. (d) PasE does not affect PerA levels. PasE overexpression does not alter PerA levels. EPEC strain expressing tagged PerA and PerB (PerA-HA, PerB-FLAG, strain LA11559) and harboring a pPasE plasmid was grown in DMEM to an OD 0.2, and expression was induced after the first hour of growth with 0.1 mM IPTG. Western blot analysis using Anti-FLAG antibodies was used to detect PerA-HA. stain-free of total protein served as Loading Control (LC).

**Supplemental information Figure 8.**
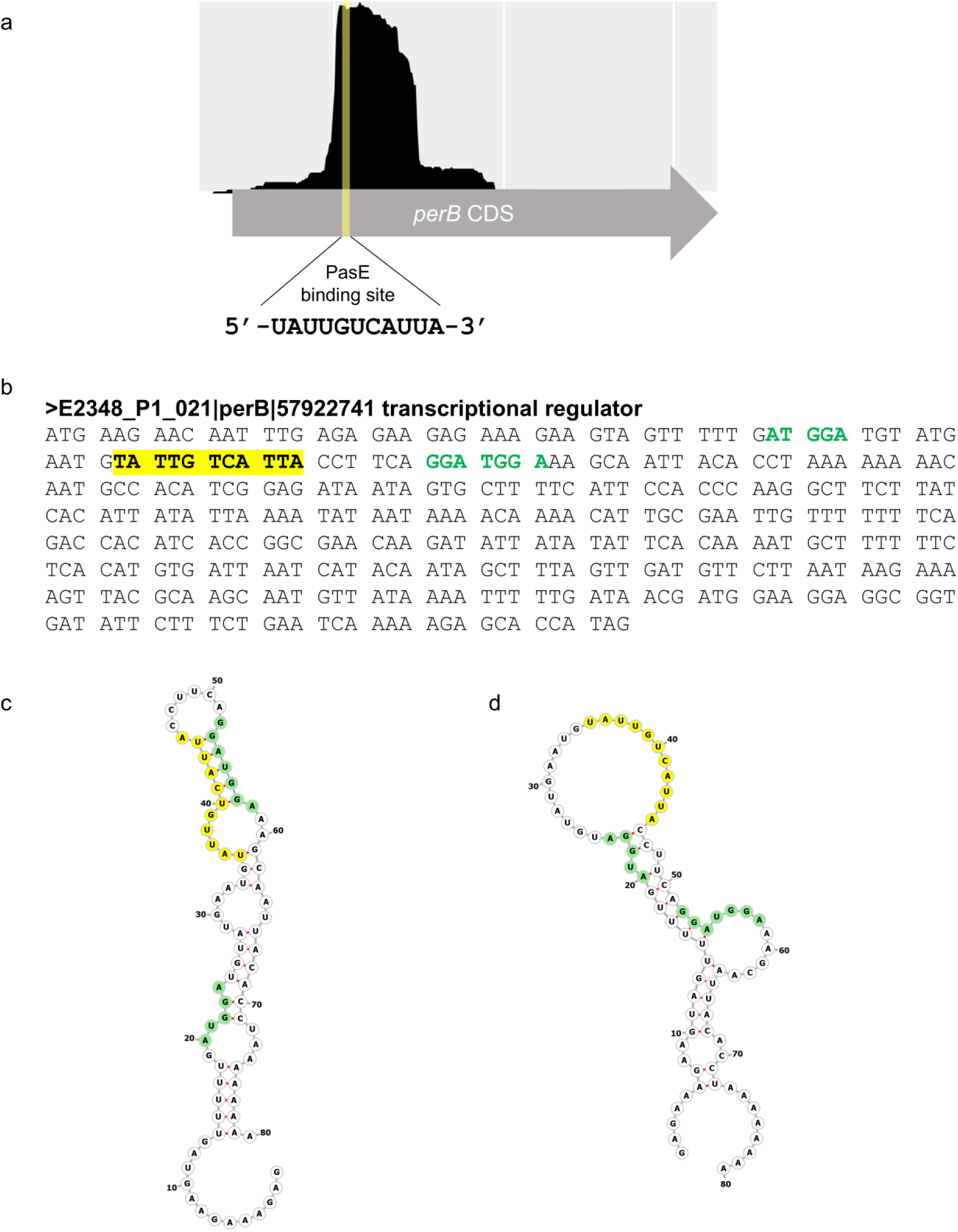
CsrA binding sites and PasE seed binding site location on the *perB* mRNA. (a) Genome browser view of the RIL-seq reads mapped to *perB* in interactions with PasE. The PasE binding site on *perB* is highlighted. (b) Nucleotide sequence of the *perB* gene, indicating the PasE seed binding site (yellow) and putative CsrA-binding GGA motifs (green). (c) Predicted secondary structure of a segment of the *perB* mRNA that includes the sequence complementary to PasE seed (marked in yellow) and the flanking putative CsrA binding sites (marked in green). The structure was predicted using Vienna RNAfold ^46^ and visualized with Forna ^49^. (d) Same region as in (c), but with a constraint applied to prevent base pairing at the sequence complementary to PasE seed (yellow), resembling the status of this region upon base pairing with PasE. Note the predicted structures suggest that initial base pairing with PasE led to exposure of CsrA binding site at nucleotides 52-58.

**Supplemental information Figure 9.**
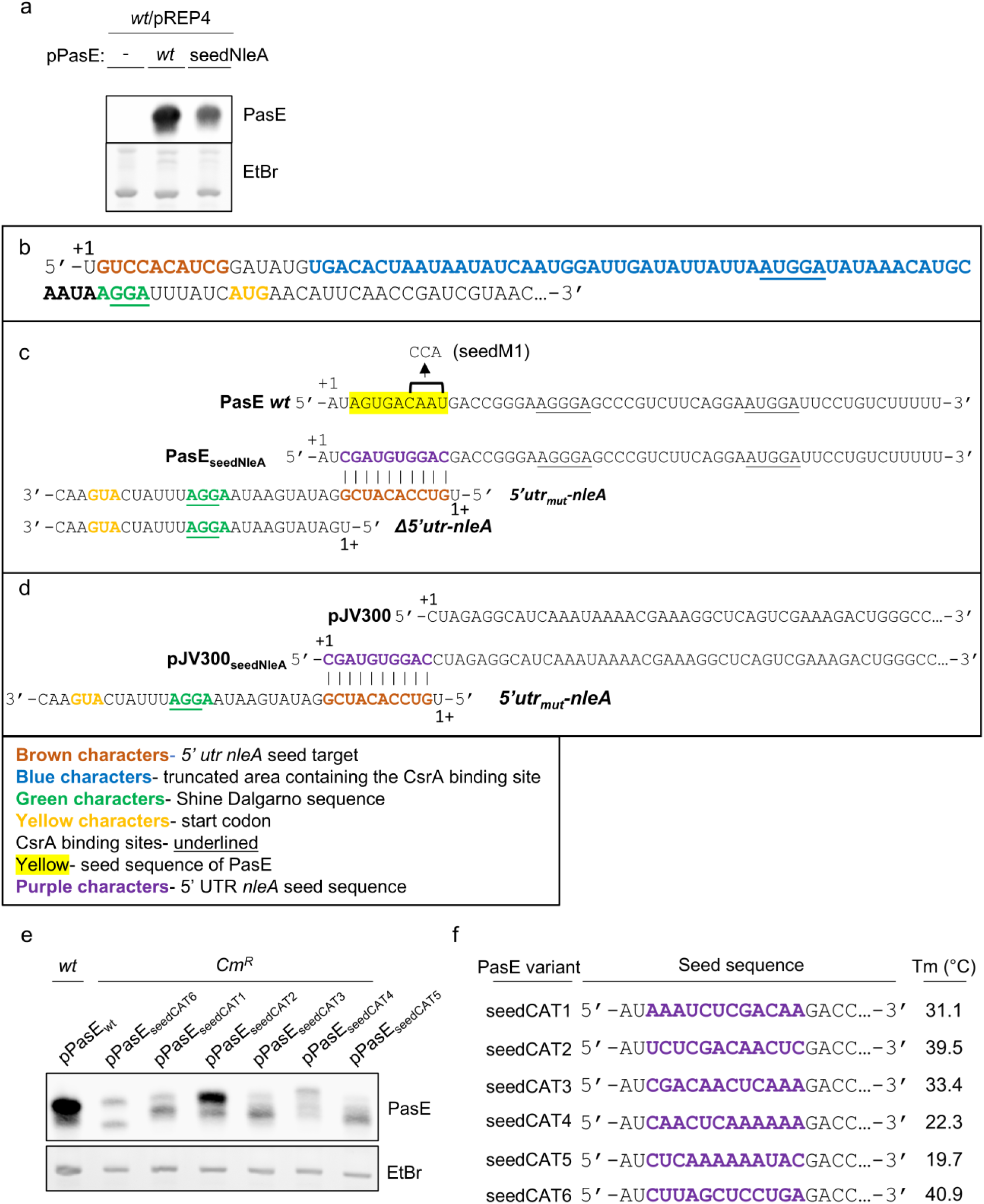
Design and analysis of PasE_seedNleA_ and JV300_seedNleA_. (a) Expression of PasE with modified seed complementing the mutated 5’ UTR of *nleA* lacking the high-affinity CsrA binding sites (PasE_seedNleA_). Wild-type EPEC ectopically expressing wild-type PasE (*wt*), or PasE_seedNleA_ variant (seedNleA), or vector (-), were grown to mid-exponential phase. pREP4 plasmids were present in all strains to express the LacI repressor. Bacteria were grown to OD ∼0.6 and expression of sRNAs was induced after the first hour of growth. RNA was extracted and subjected to Northern blot analysis. EtBr was used as a loading control for total RNA. (b) 5’ UTR sequence of the *nleA* mRNA. +1 is shown, brown characters indicate the sequence complementary to the seed of PasE_seedNleA_, and blue characters indicate the deleted area containing the high-affinity CsrA binding site, generating 5’ UTR_mut_-*nleA* (see panel c). The Shine Dalgarno sequence is highlighted in green and the CsrA binding sites are underlined. Yellow characters signify the start codon. (c) Two upper rows: comparison of the PasE native seed and the modified seed of PasE_seedNleA_. Second, third and fourth rows: The PasE_seedNleA_ seed can base pair with the 5’ UTR of the truncated *nleA* mRNA transcript (illustrated in Fig. 4c), but not with *Δ5’utr-nleA*, deleted of the complementing sequence (illustrated in Fig. 4e). (d) Design of JV300_seedNleA_ sRNA and comparison with PasE. Based on the plasmid pJV300 ^35^, we constructed a plasmid pJV300_seedNleA_ that expresses a synthetic T1 terminator fused to a seed identical to that of PasE_seedNleA_ (purple characters). The formed sRNA (JV300_seedNleA_) lacks a CsrA binding site but is still able to base pair with the *5’utr_mut_-nleA* (illustrated in Fig. 4g). The color codes corresponding to all panels are shown below panel d. (e) Expression of PasE_seedCAT_ variants. EPEC chloramphenicol-resistant strains ectopically expressing the PasE variants were grown to the exponential growth phase in DMEM. Then, RNA was extracted and analyzed by Northern blot to detect PasE. EPEC containing the pPasE plasmid was used as a reference. EtBr for total RNA staining was used as a loading control. (f) The sequence of seedCAT1-6 and their respective melting temperature, calculated using Sigma-Aldrich oligo evaluator. SeedCAT base pairing sequences are colored purple.

**Supplemental information Figure 10.**
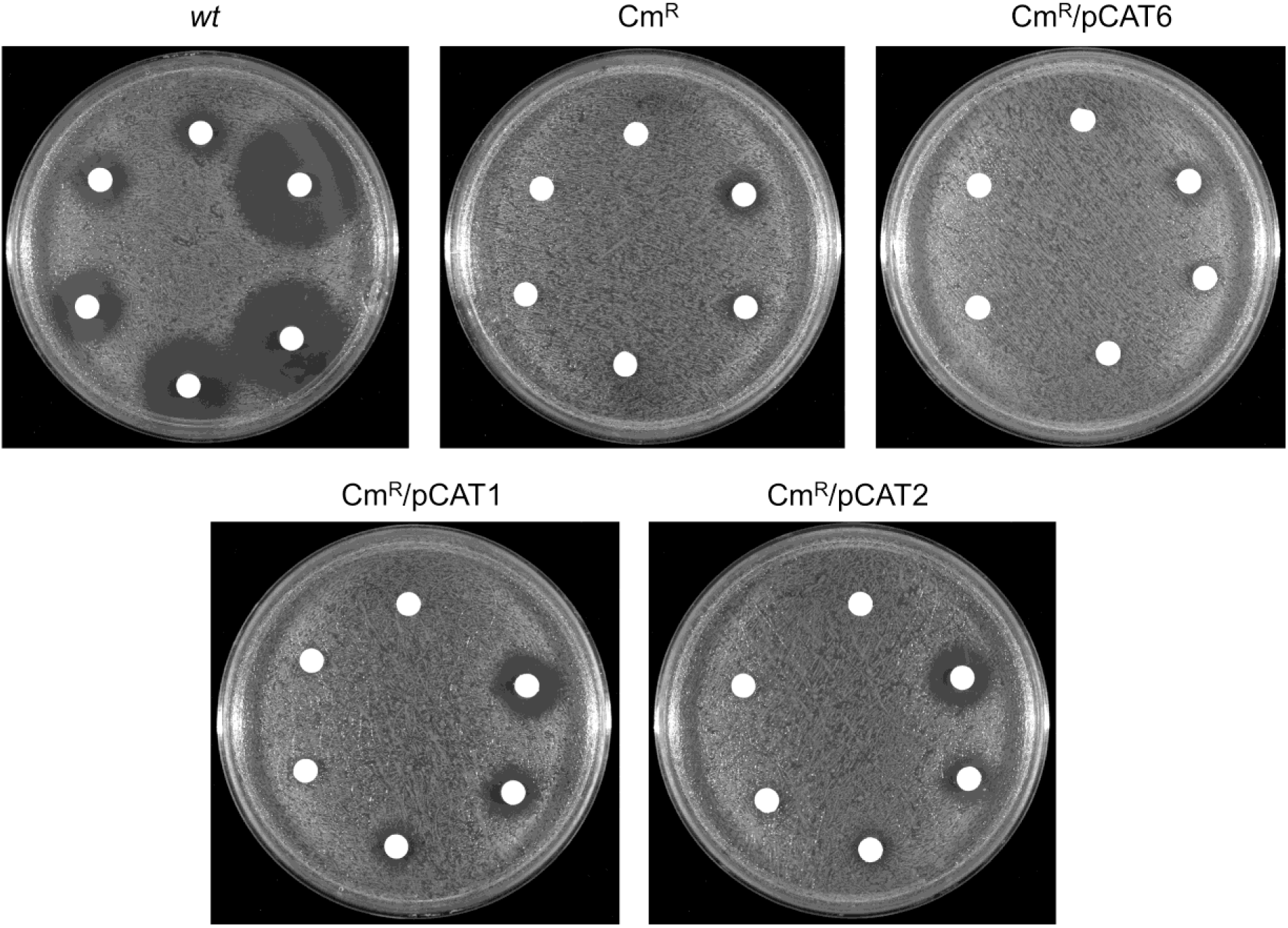
PasE_seedCAT_ variants sensitize EPEC to chloramphenicol. Wild type EPEC (*wt*) or EPEC containing a chromosomal *cat* gene and supplemented with plasmids expressing PasE variants PasE_SeedCAT1_ (Cm^R^/pCAT1), PasE_SeedCAT2_ (Cm^R^/pCAT2), PasE_SeedCAT6_ (Cm^R^/pCAT6), or remain untransformed (Cm^R^), were subjected to disk diffusion assays. Disks containing a serial dilution of chloramphenicol (Cm; 2-64 µg disk (top, lowest concentration and increasing counterclockwise) were used. Expression of PasE_SeedCAT_ variants increased bacterial sensitivity to chloramphenicol, as indicated by enlarged zones of inhibition.

**Supplementary information Figure 11.**
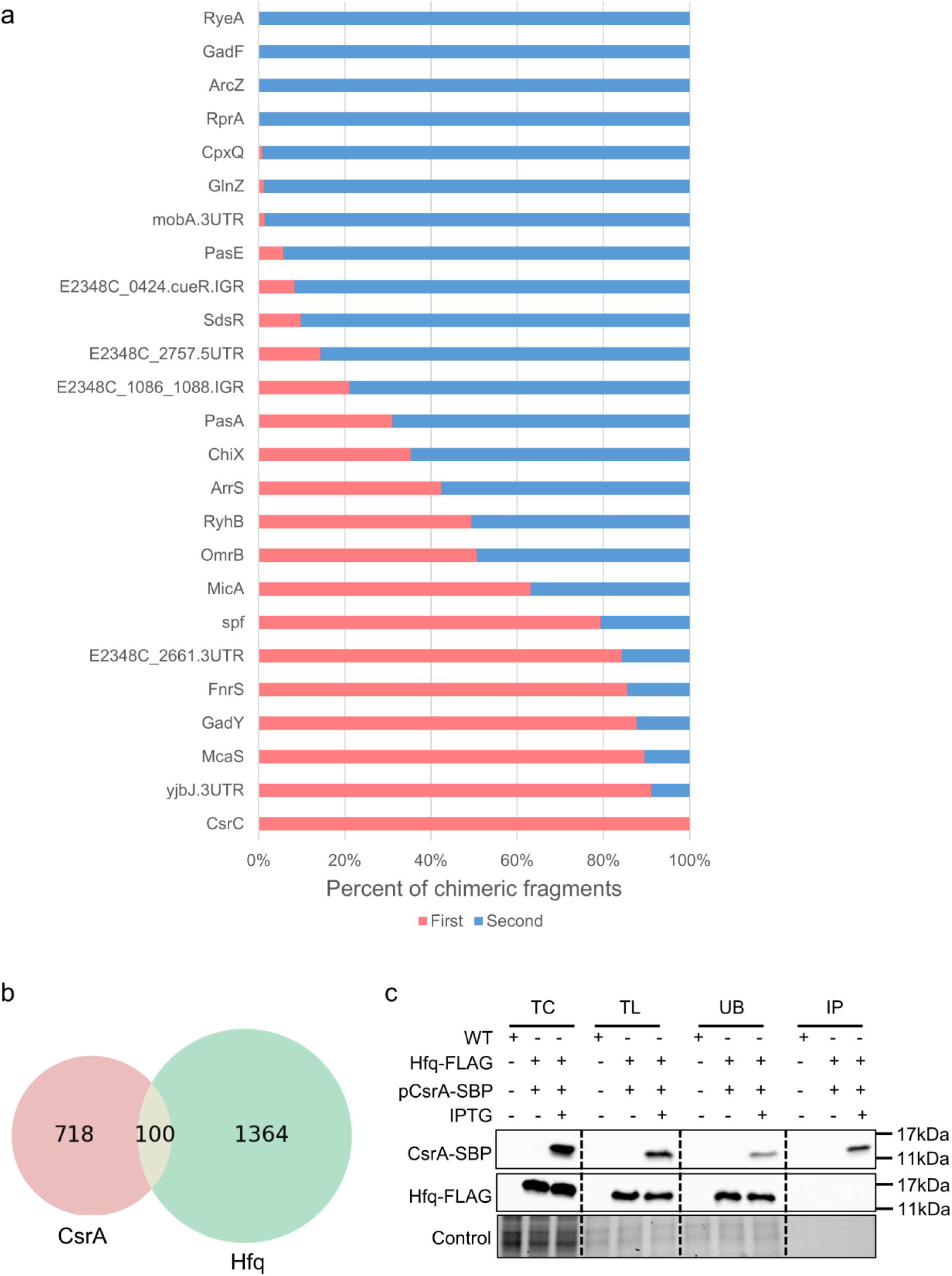
Positional preferences of sRNAs in CsrA-associated chimeras and evidence that CsrA and Hfq form distinct, non-overlapping RNA interaction networks. (a) Distribution of sRNAs by their locations at the 5’ and 3’ ends of the chimeric RNA fragments. For each sRNA found to form a chimeric fragment with > 4 different RNA molecules, we determined the percentage of the reads corresponding to it at the 5’ (First) or 3’ (Second) end of the chimeric fragments. The name of the RNA annotated as E2348C_1086.E2348C_1088.IGR in Supplementary Table 2 was shortened for convenience to E2348C_1086_1088.IGR. Interestingly, while some sRNAs equally resided in the 3’ or 5’ of the chimeric fragment (*e.g*., RyhB, OmrB), other sRNAs showed a strong preference to occupy either the 5’ (*e.g*., CsrC, *yjbJ*.3UTR) or 3’ (RyeE, GadF, PasE) of the chimeric fragment, suggesting a defined spatial organization in these interactions. (b) CsrA and Hfq bind distinct sets of RNA-RNA pairs. A Venn diagram comparing the CsrA RIL-seq (this study) data with that of the published Hfq RIL-seq data^23^. We identified only 100 RNA-RNA pairs bound to both Hfq and CsrA. The Hfq RIL-seq data included interactions from both tested conditions (activating and non-activating). This comparison revealed that CsrA and Hfq bind largely distinct sets of RNA pairs, with a minor overlap. (c) Co-immunoprecipitation experiment revealed the absence of CsrA-Hfq complex under the RIL-seq experimental conditions. EPEC *hfq-flag* (SPM6777) transformed with an inducible CsrA-SBP (Streptavidin Binding Peptide) plasmid (NE9104) was cultured in DMEM to OD_600_ of 0.2. Subsequently, CsrA-SBP expression was induced by IPTG for 2 hours until the OD_600_ reached 0.5. Bacteria were then lysed, and CsrA-SBP was immunoprecipitated from the crude extract using beads decorated with anti-SBP antibody. The beads were thoroughly washed, and the pulled proteins were analyzed using western blotting with anti-FLAG (Hfq) and anti-SBP (CsrA) antibodies. Stain-free gel of total protein served as the loading control. A wild-type EPEC strain was used as the background control. Key abbreviations: TC - Total Cells protein, TL - Total Lysate input, UB - Unbound fraction, and IP - Immunoprecipitated fraction. Notably, like in the RIL-seq protocol, we included a step of treatment with RNase and thus, in both cases, we did not expect to identify complexes of Hfq and CsrA bridged by RNAs. Taken together, the results in (B) and (C) indicate that CsrA and Hfq each cooperate with sRNAs to form two separate regulatory networks that function in parallel.

### Strains used in this study. The numbers in Name category indicate the identifier in our strain collection

| Name | Description | Source |
| --- | --- | --- |
| E2348/69 | EPEC wild-type isolate O127:H6 | J. Kaper |
| MC1061 $\lambda$ pir #2312 | *E. coli*, $\Delta(araA-leu)7697$, *araD139*, $\Delta(codB-lac)3=\Delta lac74$, *galK16*, *galE15*, *mcrA0*, *relA1*, *rpsL150*, *spoT1*, *mcrB1*, *hsdR2*, $\lambda pir$, $\lambda^-$, *F^-* | Lab collection |
| XTL634 | Template for *tet-sacB* cassette | 43 |
| #5898 | EPEC containing a Cm cassette | Lab collection |
| NN6861 | EPEC expressing a chromosomally FLAGx3 tagged CsrA in its C terminus. | This study |
| SPM6777 | EPEC expressing a chromosomally FLAGx3 tagged Hfq in its C terminus. | 23 |
| LA9372 | EPEC *perB-flagx3* | This study |
| LA9595 | EPEC $\Delta pasE$ | This study |
| LA11559 | EPEC *perA-ha*, *perB-flagx3* | This study |
| LA9156 | EPEC *PasE<sub>seedmut1</sub>* (seed mutant GAC::CCA) | This study |
| LA9543 | EPEC *perB-flagx3<sub>seedmut2</sub>* with a synonymous mutation in its target sequence (TCA::AGT) | This study |
| NE7130 | EPEC *5'utr<sub>mut</sub>-nleA* | 13 |
| LA12598 | EPEC $\Delta 5'utr-nleA$ | This study |
| NN6490 | EPEC $\Delta glgCAP::Cm^R$ | |

### Plasmids used in this study

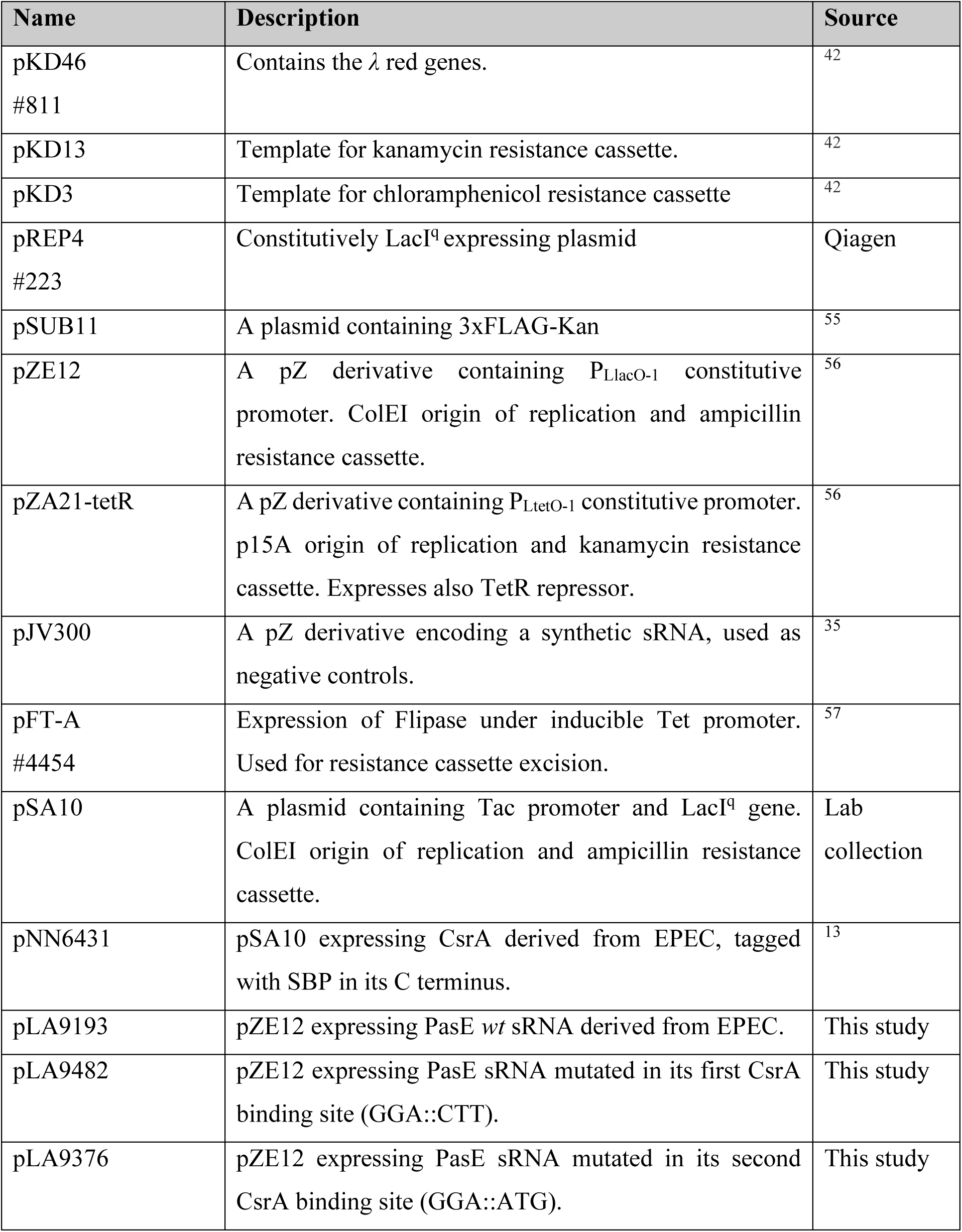
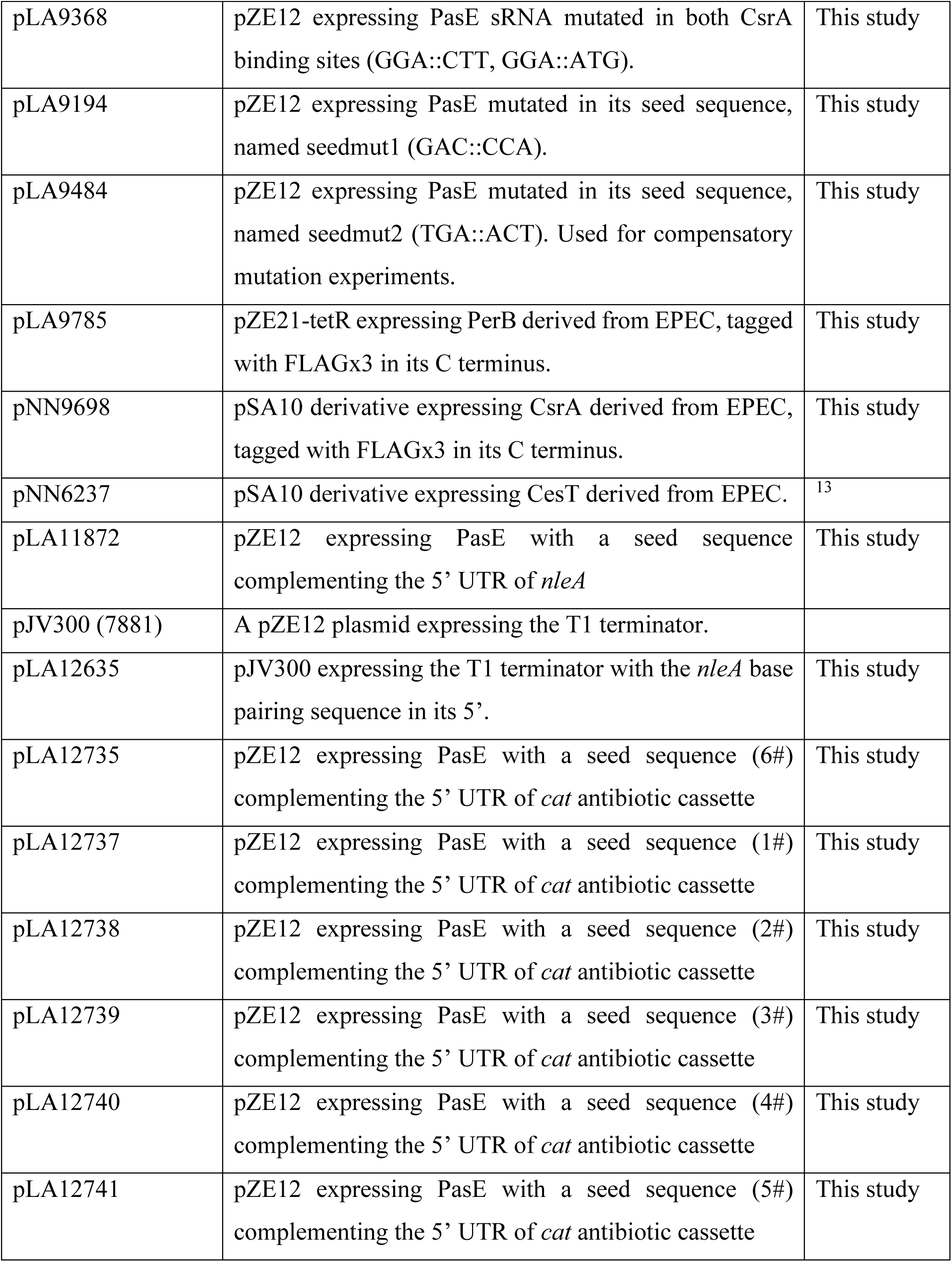

### Oligonucleotides used to construct strains

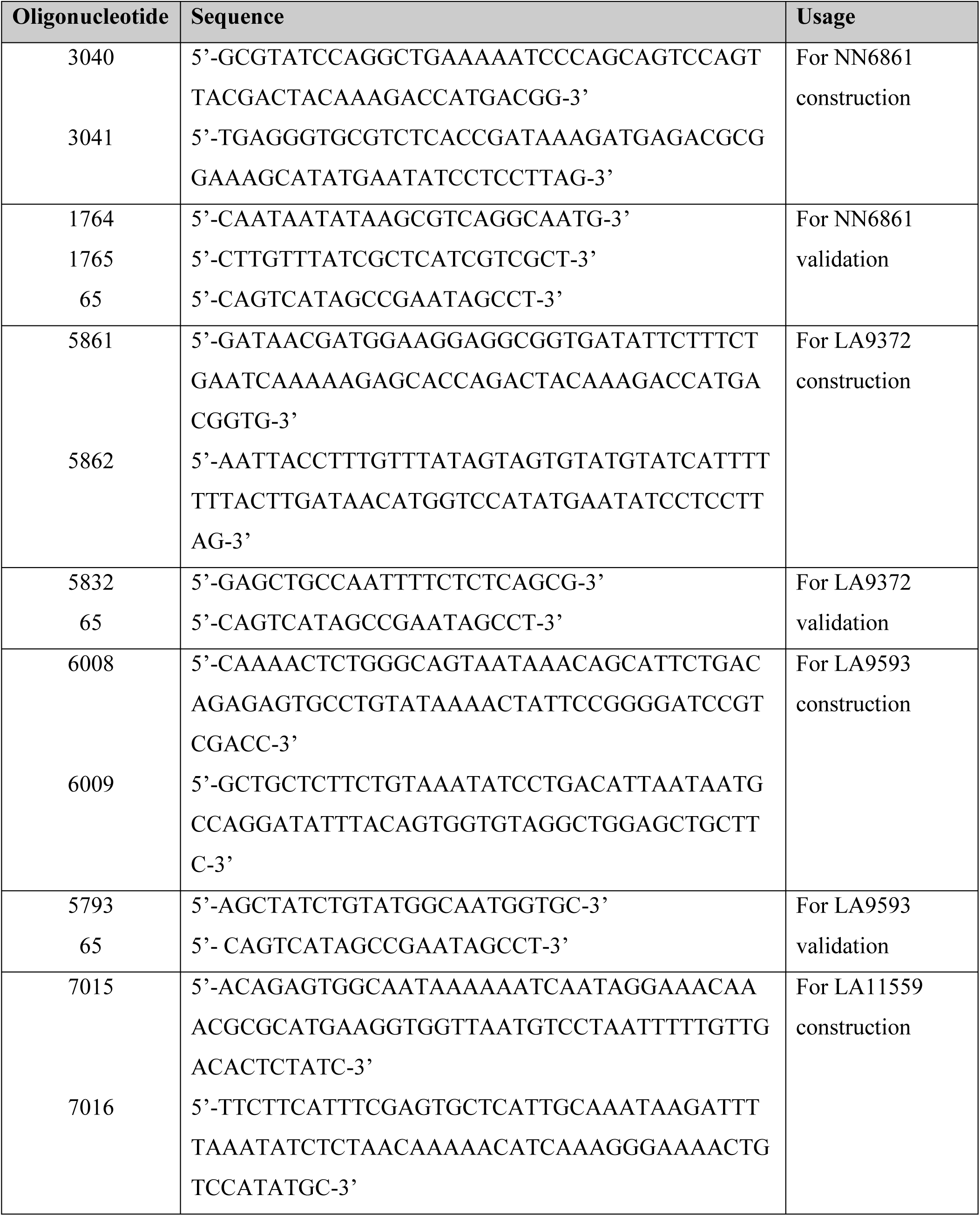
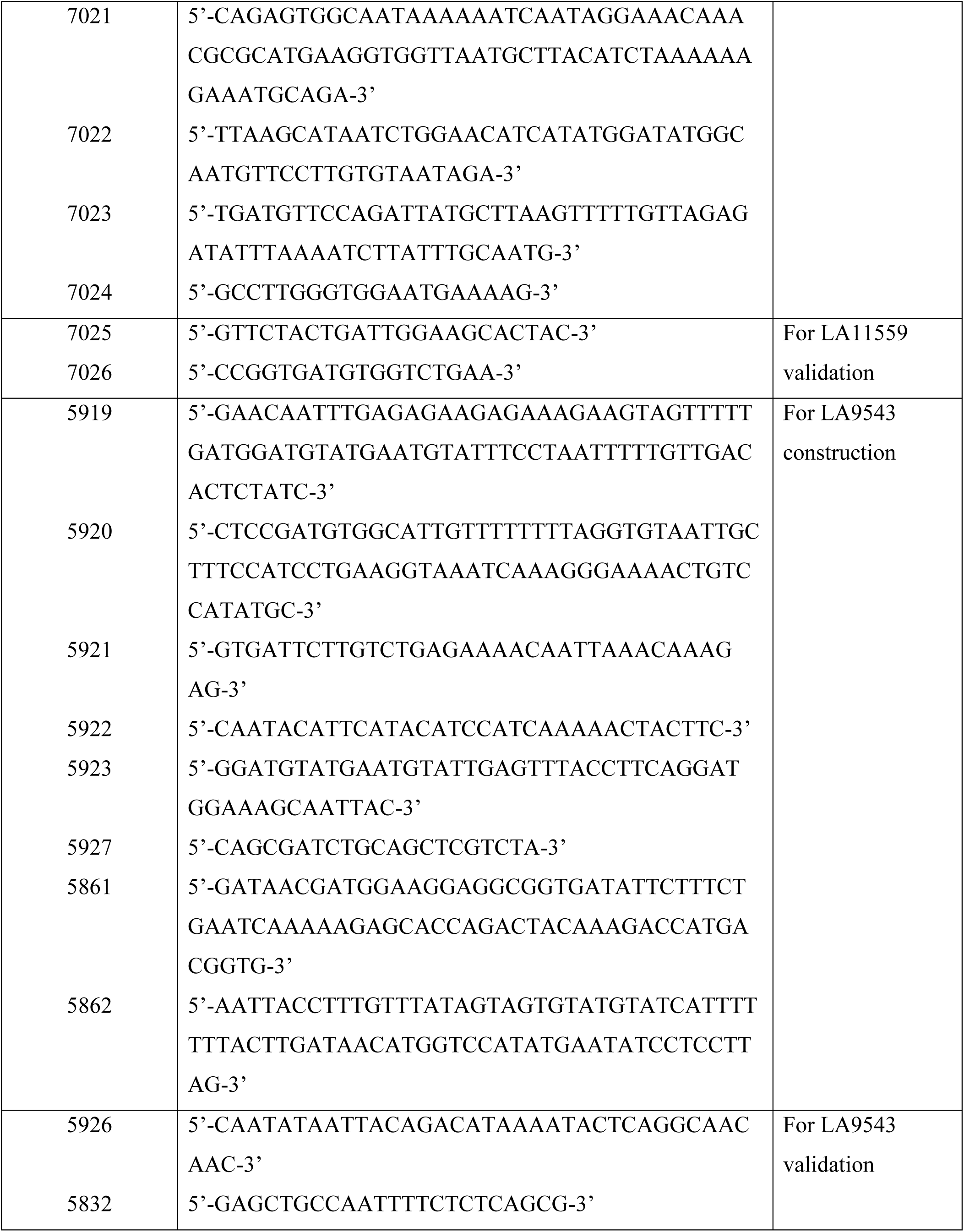
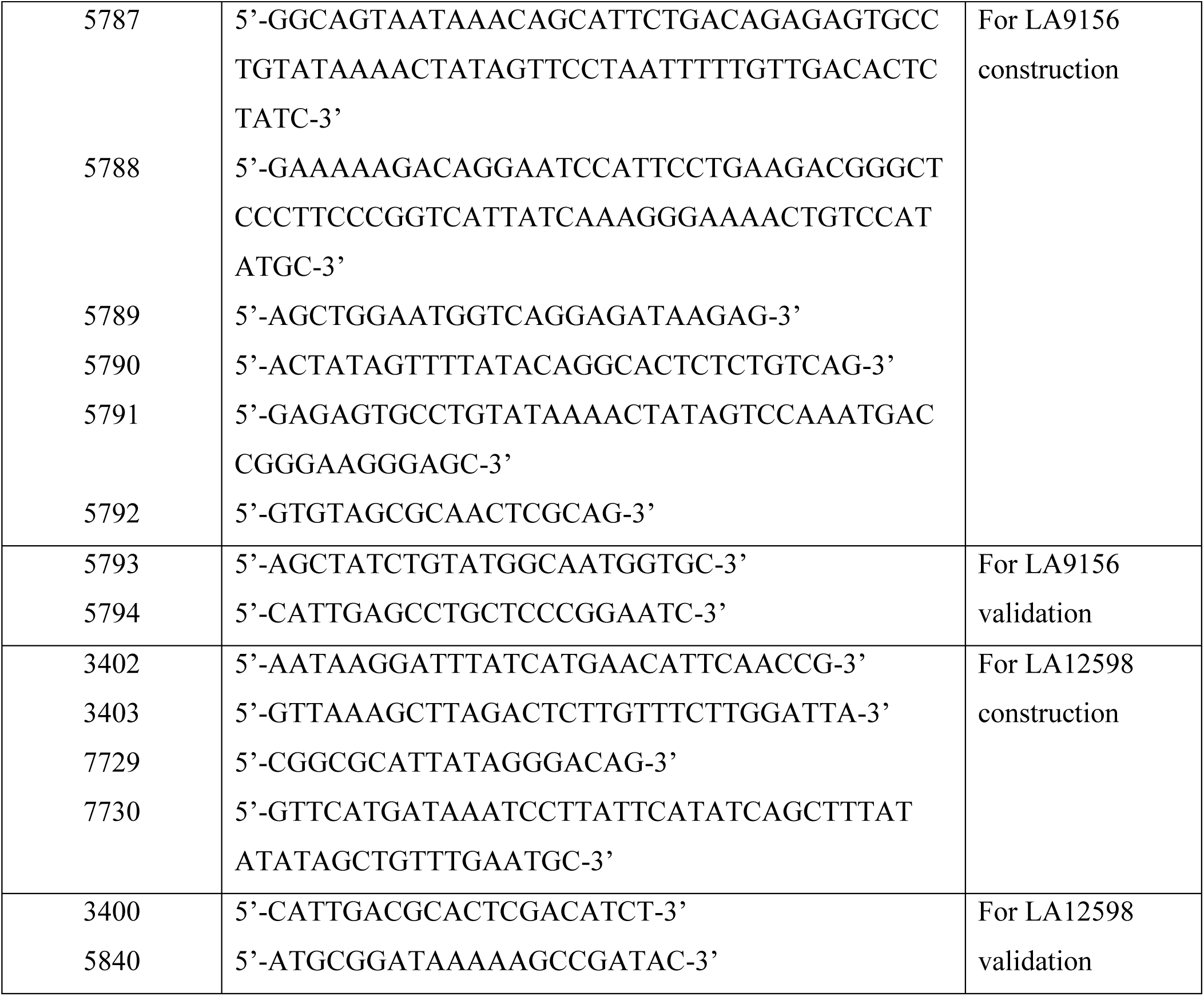

### Oligonucleotides used to construct plasmids

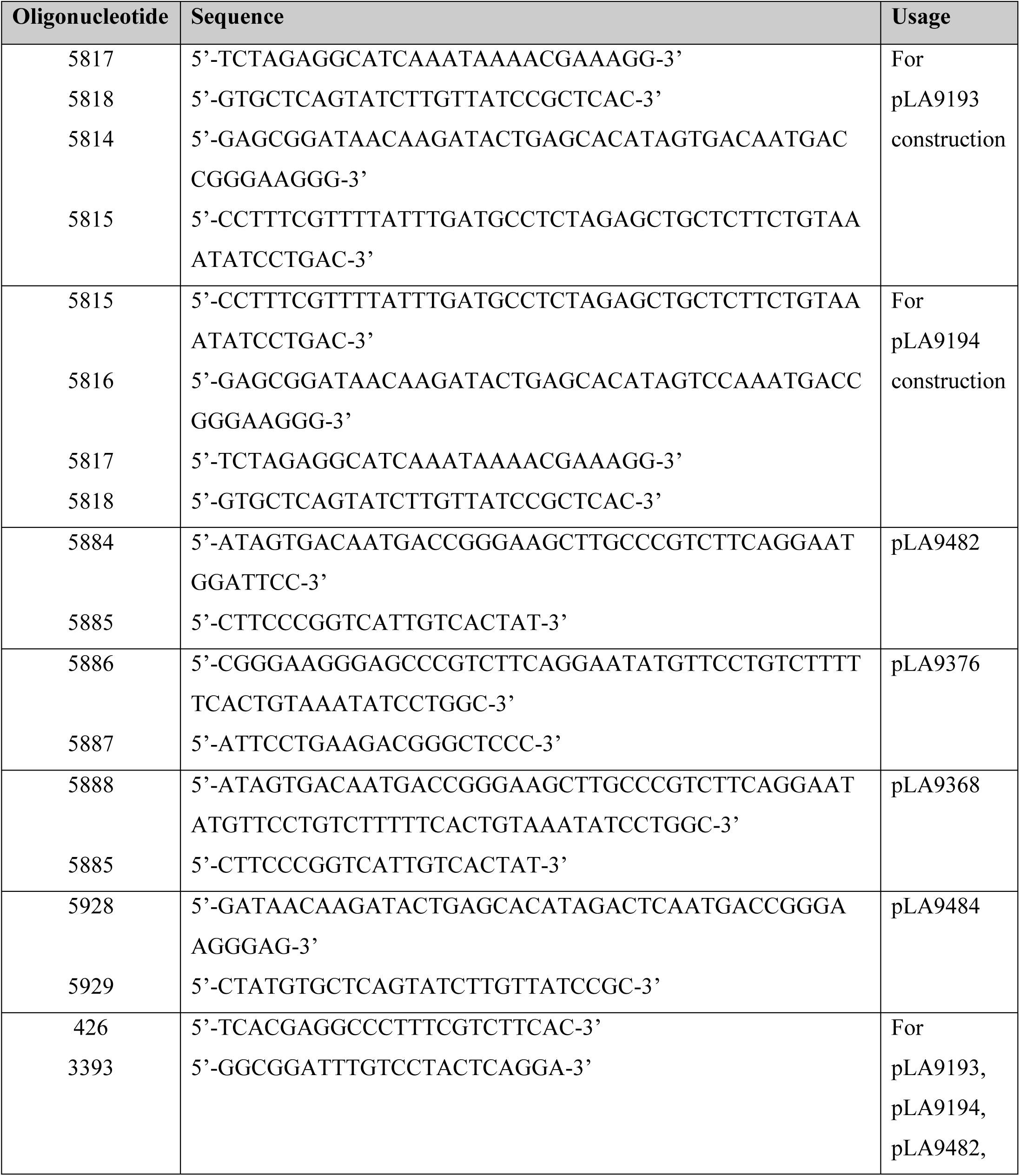
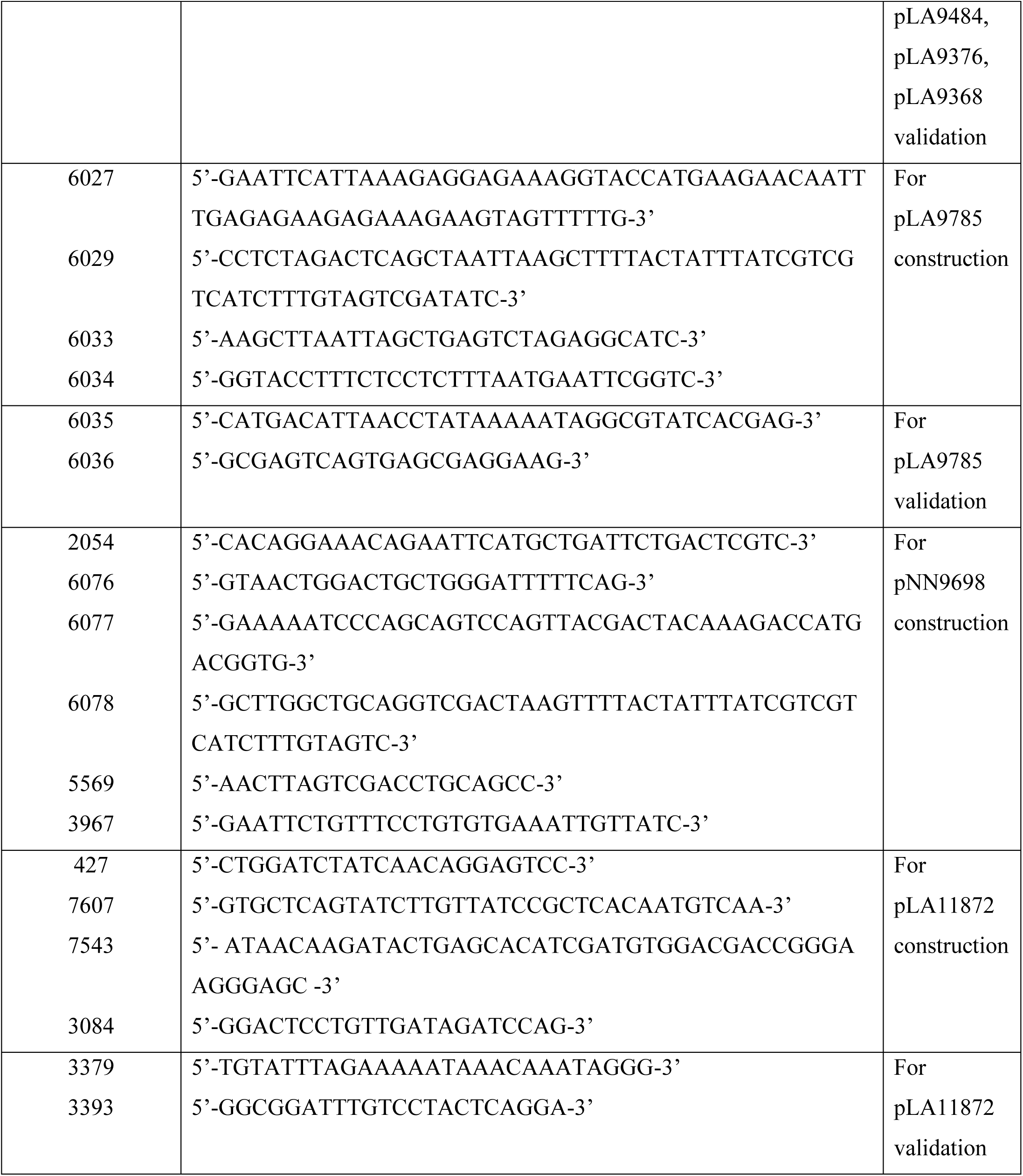
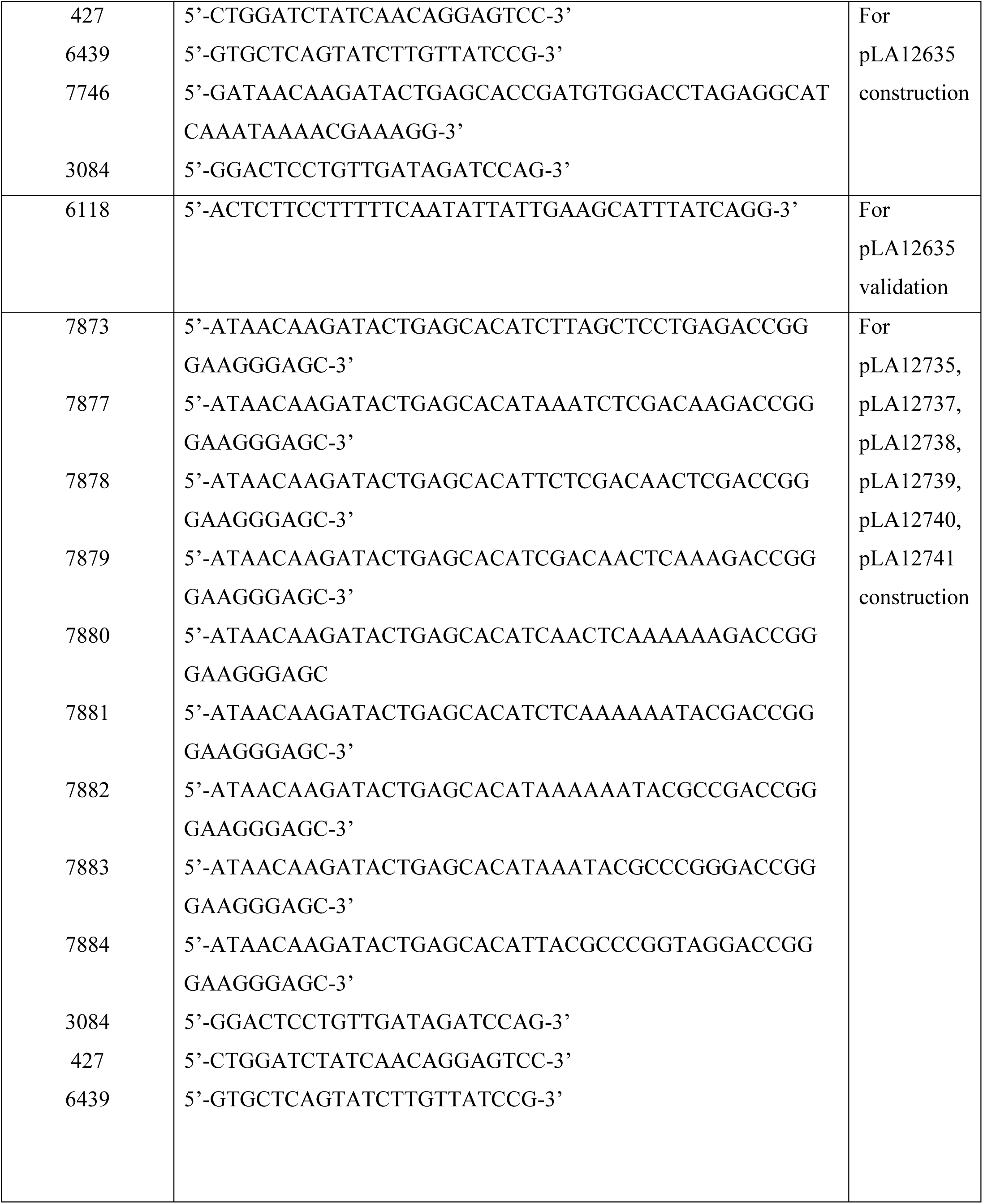
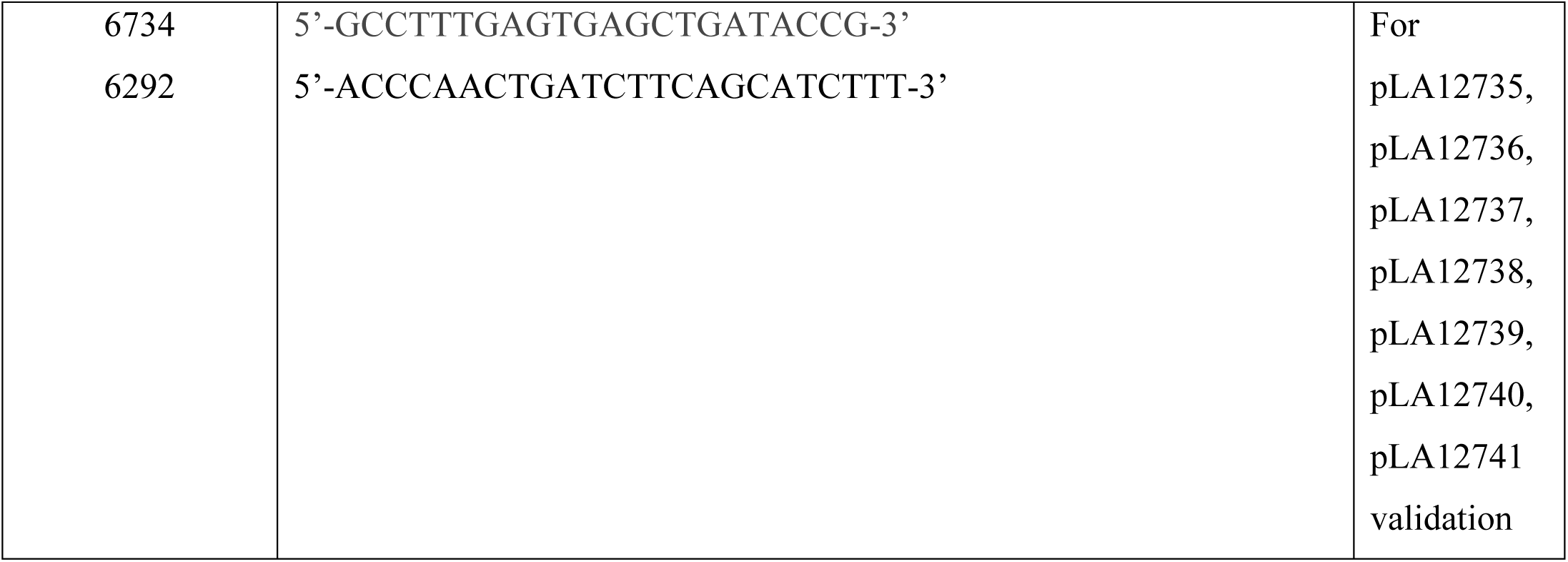

### Oligonucleotides used for RACE and northern blot experiments

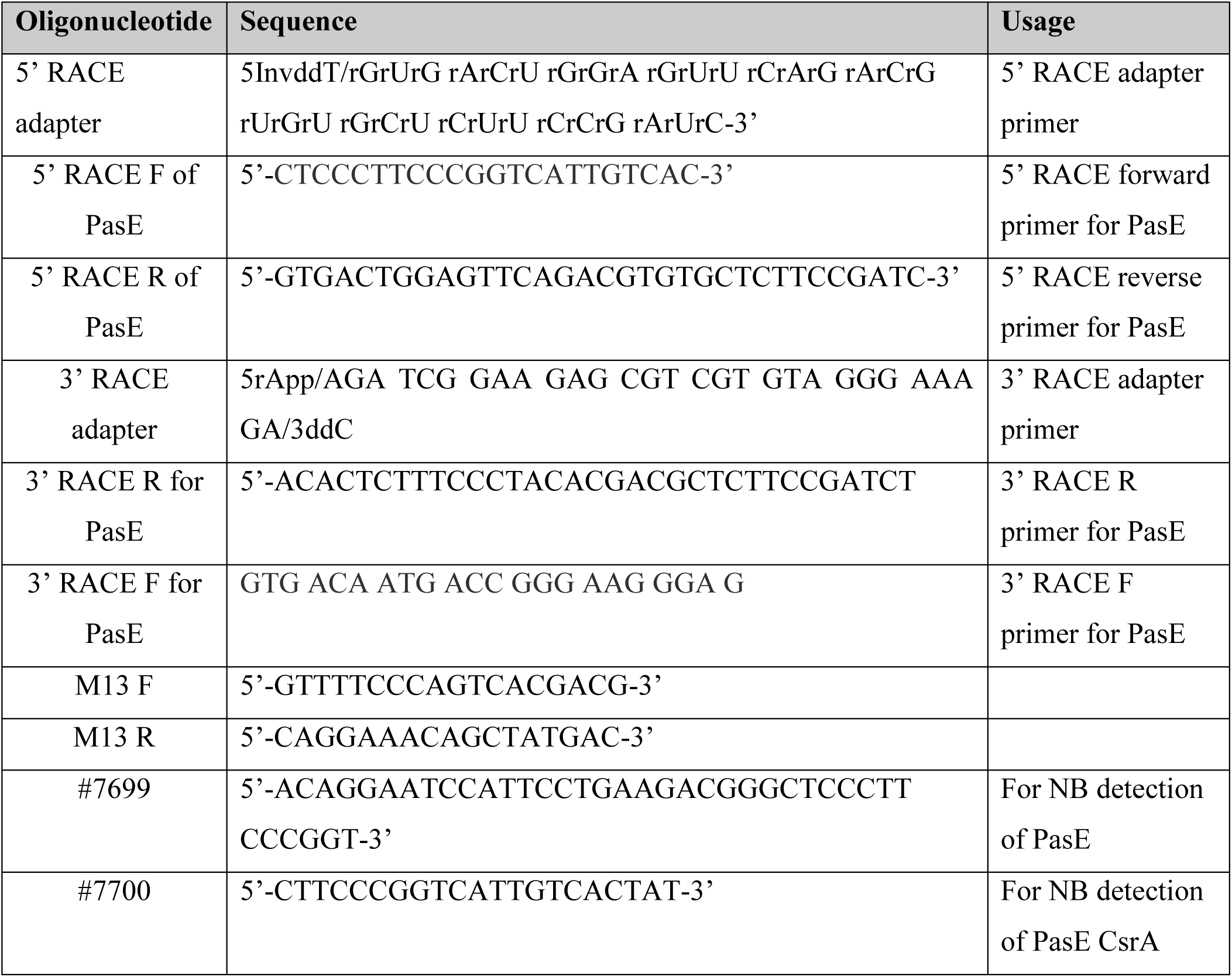
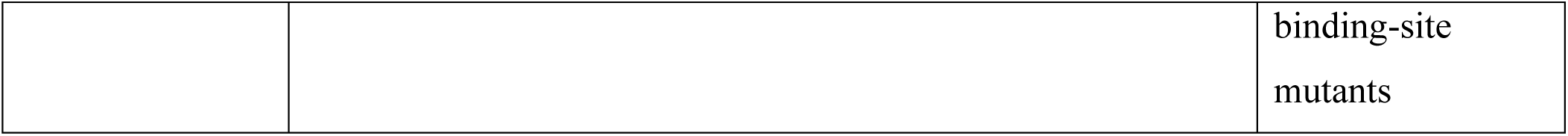

### Antibodies used in this study

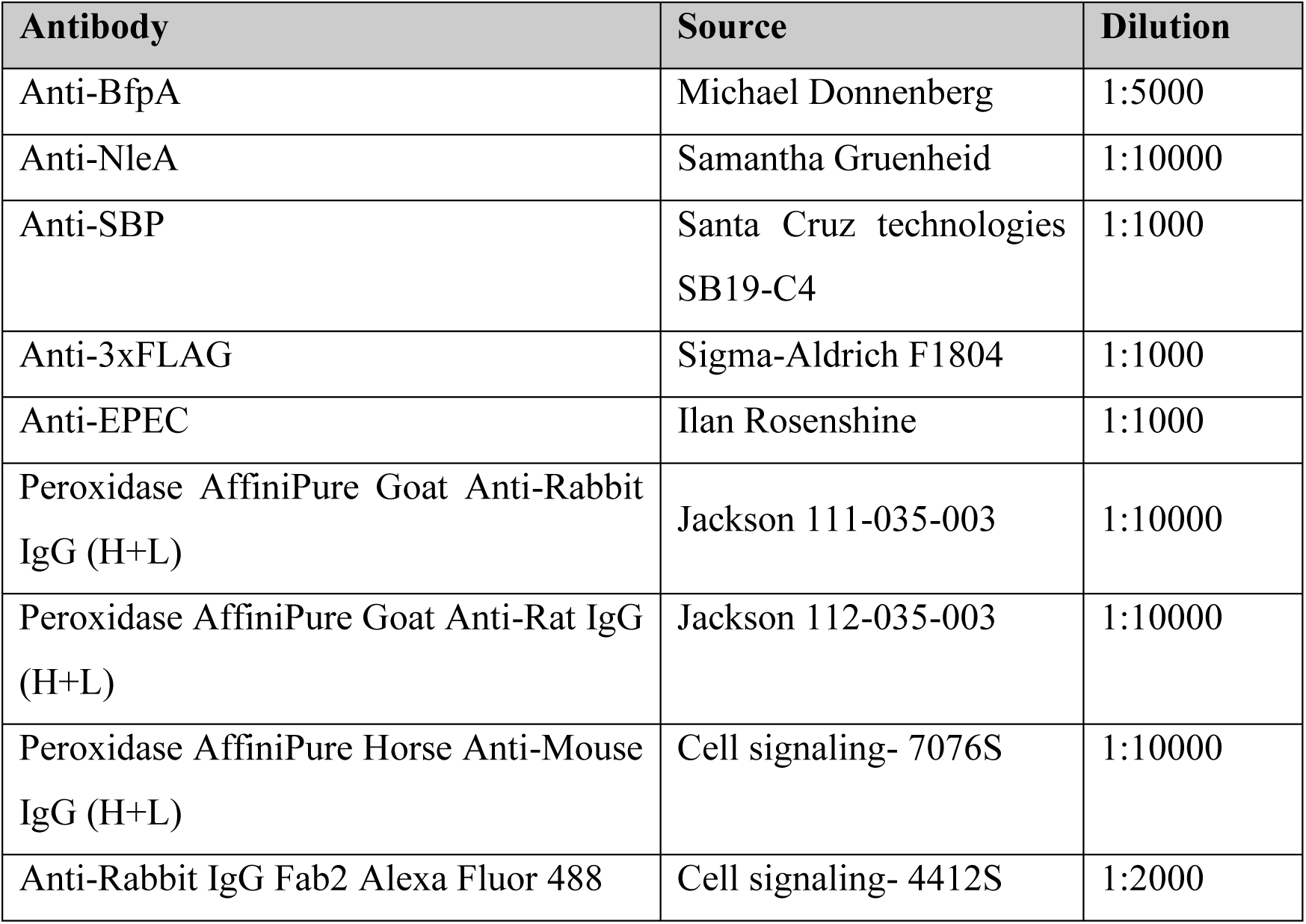

**Supplementary Table 1. Summary of number of fragments in RIL-seq sequencing libraries.** The table describes the libraries used in the experiment and statistics regarding the number of sequence fragments. The RIL-seq computational pipeline was applied to each library individually, as well as to the unified library.

**Supplementary Table 2. RIL-seq RNA pairs identified in unified datasets.**

The table includes all interactions between two RNAs in the unified dataset (minimal number of interactions ≥ 10). A pair of RNAs might appear more than once if it involves multiple interacting regions or if it appears in the chimera once as RNA1-RNA2 and once as RNA2-RNA1. Coordinates are based on the genome of EPEC E2348/69 genome version 19 (chromosome NC_011601.1 and three plasmids NC_011602.1, NC_011603.1, and EU580135). **Name**: Common name of the gene. **RNA Type**: the type of the RNA by location: 5′ untranslated region 5UTR (5′ UTR), coding sequence (CDS), 3’ untranslated region 3UTR (3′ UTR), tRNA, sRNA, non-coding RNAs (ncRNA), AS (antisense), IGR (intergenic region), and IGT (intergenic within transcript). **Number of libraries**: Number of individual libraries where this interaction was revealed as statistically significant. “0” denotes an interaction that was identified only in a unified library. **Number of chimeric fragments**: Number of chimeras supporting the interaction. **Odds Ratio**: (K/L)/(M/N), where K= Number of chimeric fragments of RNA1-RNA2, L=number of other fragments involving RNA2, M=number of other fragments involving RNA1, N=number of all other fragments (that do not involve RNA1 and RNA2). **Fisher’s exact test p-value**: p-value for observing at least this number of chimeric fragments given their background frequencies on CsrA. **Start of RNA1 first read**: Position of the first nucleotide of the most 5’ chimera mapped to the first RNA. **Start of RNA1 last read:** Position of the first nucleotide of the most 3’ chimera mapped to the first RNA. **Start of RNA2 last read:** Position of the last nucleotide of the most 5’ chimera mapped to the second RNA. **Start of RNA2 first read:** Position of the last nucleotide of the most 3’ chimera mapped to the second RNA. **Strand**: The genome strand the sequence was mapped to. **Other fragments of RNA1**: Number of fragments in which the first RNA appears as first, including single fragments. **Other fragments of RNA2**: Number of fragments in which the second RNA appears as second, including single fragments. **Core/Non-core**: core indicates that the gene is encoded on the main chromosome NC_011601 and non-core indicates that the gene is encoded on one of the three plasmids (NC_011602, NC_011603 (pEAF) and EU580135.1), other categories are encoded on NC_011601 and are listed according to Iguchi et al.^39^.

Note that when the RNA was mapped to a region outside a CDS, the name is followed by 5UTR or 3UTR in case it resides in an annotated UTR, EST5UTR or EST3UTR if the UTR is unknown and the interaction is 100 nt upstream or downstream the CDS, respectively. Two gene names and IGR or IGT represent a binding region located between two genes in two different transcription units (IGR) or on the same transcription unit (IGT). AS stands for RNA mapped to the antisense of a gene.

### Summary tab. Unique interaction

In the summary tab, each pair appears only once (disregarding the order in chimeras and regarding 5’UTR and CDS of a target as one genomic entity), where all its interactions were summed. For interactions with known sRNAs or RIL-seq putative sRNAs the sRNA was placed as RNA2, otherwise RNA1 and RNA2 were ordered alphabetically. “0” in the ‘Libraries’ columns denotes an interaction that was identified only in a unified library.

**RNA identified in CsrA singles?** “1” if the RNA was determined also as a single RNA on CsrA, according to Supplementary Table S3. **Stringent chimeras included in Figure 2**-“1” if the interaction was determined as stringent (identified in four or more libraries and was included in the network presented in Figure 2). **RNA GGA position-** the GGA sequence first coordinate is indicated. In case there were several GGA sequences the coordinates were separated with a space. **RNA GGA structure** – the predicted structure for each GGA sequence is indicated, corresponding to the coordinates order in the RNA GGA position column. The probability of accessibility was computed using RNAfold ^46^.

**Supplementary Table 3. Genes enriched in CsrA singles -results of DESeq2 analysis of EPEC CsrA-FLAG vs. wild-type CsrA**

### Accession symbol

The accession number of the gene in EcoCyc database. When the RNA was mapped to a region outside a CDS, the name is followed by 5UTR or 3UTR in case it resides in an annotated UTR (in EcoCyc), EST5UTR or EST3UTR if the UTR is unknown and the interaction is 100 nt upstream or downstream the CDS (or shorter if these regions spanned another transcript or were more likely to be a UTR of the neighboring transcript), respectively. Two gene names and IGR or IGT represent a binding region located between two genes in two different transcription units (IGR) or on the same transcription unit (IGT). AS stands for RNA mapped to the antisense of a gene.

### Name

Common name of the gene. If there is no known name for this gene the Accession symbol is displayed.

### BaseMean

The mean of normalized read counts across all the libraries (CsrA-Flag and CsrA-wt).

### Log2FoldChange

The logarithm to base 2 of the change in gene read count between EPEC CsrA-Flag and wild type.

padj: Benjamini-Hochberg adjusted p-value (corrected for multiple hypothesis testing).

### Motif Position

(A/C)NGGA Motif first positions found in this single RNA. N stands for any nucleotide.

### Motif Structure

Structure type of motifs location (loop or single strand), predicted by RNAfold tool on motif region GGA with 11 nucleotides upstream and 9 nucleotides downstream to the motif.

Motifs Total Count: Motifs total count.

Distances Between the Motifs: Distance between each two close motifs.

### Product description

description of protein product. Information by Roy Chaudhuri.

**Supplementary Table 4 - *pasE* sequence conservation across the *E. coli* and *Shigella* strains**

Sheet 1 - Legend

Sheet 2 - Web blast output file description. **Description** - description of the matching sequence (hit). **Scientific Name**- the Latin binomial classification of the source organism corresponding to each database match. **Max Score**: The score of the best local alignment. **Total Score** - Sum of alignment scores over all local alignments. **Max score = Total score** indicates that only one local alignment was found. **Query Cover** - Percent of the query length included in the alignments with the hit - represents the percentage of the query sequence length that is aligned to the subject sequence across all high-scoring segment pairs (HSPs) in the hit. **E value** - The number of hits or alignments with the same score or a better score that are expected by random. **Per. Ident** – Percent of query nucleotides that are identical to the hit sequence. **Acc. Len** - Length of the hit sequence (here it will be the length of the genome). **Accession** - accession of the hit sequence.

Sheet 3 - Taxonomy-by-scientific-name

**Scientific Name** - as in the previous sheet. **No. of occurrences** - The sum over all the occurrences of each entry in the Scientific Name field. Table entries were sorted by this field.

